# FHOD formin and SRF promote striated muscle development through separate pathways in *C. elegans*

**DOI:** 10.1101/2020.09.21.306985

**Authors:** Curtis V. Yingling, David Pruyne

**Author notes:** **Abbreviations:** body wall muscle (BWM), dilated cardiomyopathy (DCM), hypertrophic cardiomyopathy (HCM), filamentous actin (F-actin), formin homology-1 (FH1), formin homology-2 (FH2), formin homology 2 domain-containing (FHOD), myocardin-related transcription factor (MRTF), serum response factor (SRF), ubiquitin proteasome system (UPS).

## Abstract

Previous work with cultured cells has shown transcription of muscle genes by serum response factor (SRF) can be stimulated by actin polymerization driven by proteins of the formin family. However, it is not clear if endogenous formins similarly promote SRF-dependent transcription during muscle development *in vivo*. We tested whether formin activity promotes SRF-dependent transcription in striated muscle in the simple animal model, *Caenorhabditis elegans*. Our lab has shown FHOD-1 is the only formin that directly promotes sarcomere formation in the worm’s striated muscle. We show here FHOD-1 and SRF homolog UNC-120 both support muscle growth and also muscle myosin II heavy chain expression. However, while a hypomorphic *unc-120* allele blunts transcription of a set of striated muscle genes, these genes are upregulated or unchanged by absence of FHOD-1. Instead, pharmacological inhibition of the proteasome restores myosin protein levels in worms lacking FHOD-1, suggesting elevated proteolysis accounts for their myosin deficit. Interestingly, proteasome inhibition does not restore normal muscle growth to *fhod-1(Δ)* mutants, suggesting formin contributes to muscle growth by some alternative mechanism. Overall, we find SRF does not depend on formin to promote muscle gene transcription in a simple *in vivo* system.

## Introduction

Dilated and hypertrophic cardiomyopathies, pathological thinning or thickening of the ventricle wall of the heart, respectively, correlate with a dysregulation of cardiac muscle growth [1]. Dilated cardiomyopathy (DCM) occurs in ∼1 in 2500 individuals, while hypertrophic cardiomyopathy (HCM) occurs in ∼1 in 500 individuals [1]. Significantly, in the United States, HCM is the leading cause of sudden cardiac-related death of young competitive athletes [2]. Many of the genetic causes of DCM and HCM have been linked to mutations in components of the sarcomere [1].

The sarcomere is the basic unit of contraction in muscle cells. Thick filaments are the contractile component of the sarcomere, whose myosin motors pull on thin filaments composed of actin and associated proteins. In vertebrate striated muscle, thin filaments are in turn attached to Z-discs that define the ends of the sarcomere. Two well-studied examples of genetic causes of DCM and HCM are β cardiac myosin heavy chain (encoded by *MYH7*) of the thick filament, and cardiac troponin I (encoded by *TNNI3*) of the thin filament [1]. Another sarcomeric component of cardiac muscle is formin homology 2 domain containing 3 (FHOD3) [3, 4], a member of the formin family of proteins. Mutations in the *FHOD3* gene have recently been shown to be a genetic cause of HCM, while other mutations have been associated with DCM [5–11].

Formins in general are actin nucleation factors that have the unique ability to also act as “leaky” or “processive” caps that protect the growing barbed end of actin filaments from inhibitors [12–17]. Many formins, including FHOD3, exist in an autoinhibited state where intramolecular interactions inhibit activity, and are activated by regulated disruption of these intramolecular interactions [18]. Several lines of evidence demonstrate FHOD3 plays a fundamental role in sarcomere organization in cardiac muscle. In cultured neonatal rat cardiomyocytes, loss of FHOD3 leads to disrupted sarcomere formation, while expression of mutationally hyperactivated FHOD3 leads to an increase in cell surface area [19, 20]. In mice, knock out of the *FHOD3* gene leads to death from heart failure by embryonic day 11.5 with appearance of immature sarcomere-containing premyofibrils/stress fiber-like structures with immature Z-lines that fail to mature into myofibrils [21]. In contrast to the full knockout or knockdowns of *FHOD3* in mouse or cultured cell studies, most identified human variants of *FHOD3* related to HCM or DCM are missense mutations [5–11].

Interestingly, formins in general and FHOD3 in particular have been implicated in the proper function of serum response factor (SRF), another protein thought to be involved in DCM and HCM. Rather than being a sarcomere component, SRF is a ubiquitous transcription factor [22]. Cardiac-specific inactivation of SRF using β myosin heavy chain-driven CRE in mice, leads to lethal cardiac defects in utero [23], and cardiac-specific SRF inactivation induced during adulthood in mice leads to DCM and death within 10 weeks, whereas forced overexpression of SRF in mouse heart leads to cardiac hypertrophy [24, 25]. Human variants in the *SRF* gene have not been directly linked to DCM or HCM, but dominant negative splice isoforms and caspase 3-cleavage products of SRF are expressed in failing human hearts [26, 27]. These data collectively suggest alteration of SRF activity may be involved in heart disease.

Alone, SRF is a weak transcriptional activator that relies on over 60 cofactors to alter its transcriptional activity in different contexts [28]. Myocardin, the founding member of the myocardin-related transcription factor (MRTF) family, is a strong activating cofactor primarily expressed in the cardiovascular system [22]. Myocardin causes SRF to selectively transcribe smooth muscle- and cardiac muscle-specific genes, such as those encoding smooth muscle myosin heavy chain and cardiac-specific α and β myosin heavy chains [29, 30]. Related family members MRTF-A and MRTF-B are also strong activating cofactors of SRF [31], but while myocardin is constitutively nuclear, MRTF-A and MRTF-B subcellular distributions are regulated by the actin monomer concentration in the cell. That is, actin monomers sequester MRTF-A and MRTF-B in the cytoplasm, whereas actin polymerization will reduce monomer concentration and release MRTFs to enter the nucleus and bind SRF to drive smooth muscle gene transcription [32].

As proteins that promote actin polymerization, mammalian formins, including FHOD3, have been shown to drive MRTF-dependent transcription of SRF-dependent luciferase reporter constructs, particularly when the formins are expressed as activated constructs [33–38]. Interestingly, FHOD3 Y1249N, a human *FHOD3* variant identified in DCM patients, has reduced ability to stimulate SRF activity, suggesting FHOD3 Y1249N is only partially functional [5]. While these data leave open the possibility that FHOD3 and SRF/MRTF work through the same pathway to affect progression of DCM and HCM, it remains to be demonstrated whether formins activate the SRF/MRTF pathway during muscle cell development *in vivo*.

The genetically tractable roundworm *Caenorhabditis elegans* provides a simple *in vivo* model to test whether SRF and formins coordinate in striated muscle development. The largest muscle group in the worm, the striated body wall muscle (BWM), has long been a model for cardiac and skeletal muscle function, with many homologous sarcomere and other muscle proteins, including homologs for SRF (worm UNC-120) and FHOD3 (worm FHOD-1) [39]. Worm FHOD-1 is expressed in all muscle types and is the only formin that promotes BWM growth in a cell autonomous manner [40, 41]. One advantage to the use of *C. elegans* as a model system is that absence of FHOD-1 is not lethal, unlike loss of FHOD3 in mice, allowing investigation of the effects of FHOD-1 loss on striated muscle at later stages of development. A deletion in the actin-binding formin homology-2 (FH2) domain coding region of the *fhod-1* gene results in slow BWM growth due to fewer sarcomeres being assembled per BWM cell [40]. These worms lacking functional FHOD-1 also have a disrupted organization of dense bodies [42], which serve as the primary sarcomere Z-line structures that anchor thin filaments in BWM. Of significance to this study, *fhod-1* mutant worms also have a decreased expression of a muscle-specific myosin heavy chain (MYO-3) [42], as might also be expected if FHOD-1 drives SRF activity toward transcription of BWM genes, similar to its counterpart in mammalian cultured cell systems.

The worm SRF-coding gene, *unc-120*, was discovered in a forward genetic screen for mutations that cause uncoordinated movement in *C. elegans* [43]. UNC-120 is expressed in muscle cells, and its overexpression in embryonic blastomeres is sufficient to convert cells to a BWM-like cell fate, based on their novel expression of muscle-specific MYO-3 [44]. Similar BWM-enriched embryos created by RNAi-based knockdown of *mex-3*, *skn-1*, and *elt-1*, had increased expression of 2058 putative muscle genes, 40% of which depended on the presence of *unc-120* [45]. Based on these results, UNC-120 is thought to be a muscle-specific transcription factor, in contrast to mammalian SRF, which functions in multiple pathways. Worms bearing a deletion allele of *unc-120* eliminating its start codon and most of the sequences encoding DNA-binding and dimerization domains, are normal early in their first larval (L1) stage, but experience progressive paralysis thereafter, with death by early L2 stage [44]. However, worms bearing a temperature-sensitive *unc-120(st364)* mutation are viable when maintained at semi-permissive temperature, but interestingly have narrow muscle cells that superficially appear similar to those of worms lacking FHOD-1 [46].

In this paper, we examine whether FHOD-1/formin promotes UNC-120/SRF transcriptional activity to promote *in vivo* muscle cell growth and MYO-3 expression in *C. elegans*.

## Materials and Methods

### Worm strains and growth conditions

Worms were maintained on nematode growth medium (NGM) plates with OP50-1 bacterial lawns for food, and were handled using standard laboratory procedures for *C. elegans* [43]. Worms were grown at 20°C, or at 15°C for experiments using temperature-sensitive animals. Wild-type strain N2 and temperature-sensitive strain RW364 [*unc-120(st364)* I] [46] were obtained from the *Caenorhabditis* Genetics Center (University of Minnesota, Minneapolis, MN). XA8001 [*fhod-1(tm2363)* I] and DWP10 [*fhod-1(tm2363)* I; *qals8001[unc-119(+) fhod-1::gfp]*] were previously described [40]. Age-synchronized populations were obtained by one of two methods. For most experiments, gravid adults were allowed to lay eggs on OP50-1/NGM plates 4 hours before adults were removed, resulting in semi-synchronized progeny. To collect larger numbers of age-synchronized worms for mRNA sequencing, embryos were collected by washing gravid adults from OP50-1/NGM plates and treating them 25 min on ice with alkaline bleach to release embryos, which were washed in M9 medium [47] and plated onto OP50-1/NGM plates.

### Fluorescence microscopy

Worms were stained for filamentous actin (F-actin) for fluorescence microscopy, as previously described [40]. Briefly, animals were suspended in M9 and fixed by addition of equal volume phalloidin mix (160 mM KCl, 40 mM NaCl, 1 mM EGTA, 30 mM NaPIPES, pH 7.3, 100 mM spermidine-HCl, 1:125 Alexa Fluor 568-phalloidin; ThermoFisher Scientific, Waltham, MA), and 0.1 volume 20% formaldehyde, and subjected to three rapid freezes in liquid nitrogen and thaws in 65°C water bath until near completion. After the final thaw, worms were incubated 1 hour on ice for fixation, washed twice 1 min with Tris-Triton (100 mM Tris-HCl, pH 7.5, 1 mM EDTA, 1% Triton X-100), and once 15 min with PBST-B (PBS, 1 mM EDTA, 5 mM NaN_3_, 0.5% Triton X-100, 0.1% BSA). Fixed animals were mixed overnight at room temperature in PBST-B/1:250 Alexa Fluor 568-phalloidin, washed twice 25 min with PBST-B, once 25 min with 1 μg/mL 4’,6-diamidino-2-phenylindole in PBST-B, and once 25 min with PBST-B before mounting.

Immunostain of worms for fluorescence microscopy was modified from previously described [48]. Monoclonal antibody 5-6-s (anti-MYO3) generated by H.F. Epstein (Baylor College of Medicine, Houston, TX) was obtained through the Developmental Studies Hybridoma Bank (University of Iowa, Iowa City, IA), and secondary antibody Texas red-conjugated goat anti-mouse was commercially obtained (Rockland Immunochemicals, Limerick, PA). 50 μL packed worms were suspended in M9, chilled 5 min on ice, resuspended in 1 mL (80 mM KCl, 20 mM NaCl, 10 mM EGTA, 15 mM Na PIPES, pH 7.4, 75% methanol) plus 25 μL 20% formaldehyde, and immediately frozen in liquid nitrogen. Samples were subjected to five rapid freezes with liquid nitrogen and thaws in 65°C water bath until near completion, after which samples were fixed 30 min on ice. Fixed worms were washed/incubated twice 1 min with Tris-Triton, 2 hours with Tris-Triton/1% β-mercaptoethanol, 1 min with BB (25 mM H_3_BO_3_, 2.5 mM NaOH, pH > 9.5), 20 min with BB/10 mM dithiothreitol, 1 min with BB, 20 min with BB/0.3% H_2_O_2_, 1 min with BB, and 15 min with PBST-B. Samples were incubated overnight with 5-6-s in PBST-A (PBST-B with 1% BSA), washed three times 25 min with PBST-B, incubated 4 hours with secondary antibodies (that were precleared overnight against similarly fixed animals) diluted 1:500 in PBST-A, washed twice 25 min with PBST-B, once 25 min with PBST-B containing 1 μg/mL 4’,6-diamidino-2-phenylindole, and once briefly with PBST-B before mounting.

Wide-field fluorescence images were acquired on an Eclipse 90i research upright microscope (Nikon, Tokyo, Japan) at room temperature using a CFI Plan Apochromat 40X/NA 1.0 oil immersion objective or CFI Plan Apochromat violet corrected 60x/NA 1.4 oil immersion with a Cool-SNAP HA2 digital monochrome charge-coupled device camera (Photometrics, Tuscon, AZ) driven by NIS-Elements AR acquisition and analysis software (version 3.1; Nikon, Tokyo, Japan).

### Microscopy image analysis

Images for quantitative analyses in Fig. 1 and Fig. 5 A were acquired using autoexposure and brightened for easy viewing using Photoshop (Version 21.1.0; Adobe, San Jose, CA). Images in Fig. 5 F were acquired using consistent exposure times to provide a rough visual comparison of relative levels of MYO-3, and all images in the panel were linearly processed identically using Photoshop to enhance visibility. All quantitative analyses of images were performed while blinded to sample identity. Distance measurements were made using the two-point tool of NIS-Elements, and exported to Excel (Microsoft, Redmond, WA) for processing. Measured BWM widths were the lateral widths of the F-actin-rich contractile lattices of one ventral and one dorsal BWM quadrant in phalloidin-stained worms, as shown by double-arrows in Fig. 1 A. BWM cell widths in phalloidin-stained worms were measured across the widest portion of the F-actin contractile lattice of individual BWM cells, and striations per cell were determined by manually counting the visible striations in the same BWM cells. Overall body width was measured based on autofluorescence of the worm body in the fluorescein isothiocyanate channel. As BWM width and overall body width vary based on anterior/posterior position in the animal, measurements were consistently performed at a position three to five muscle cells away from the vulva.

**Figure 1.**
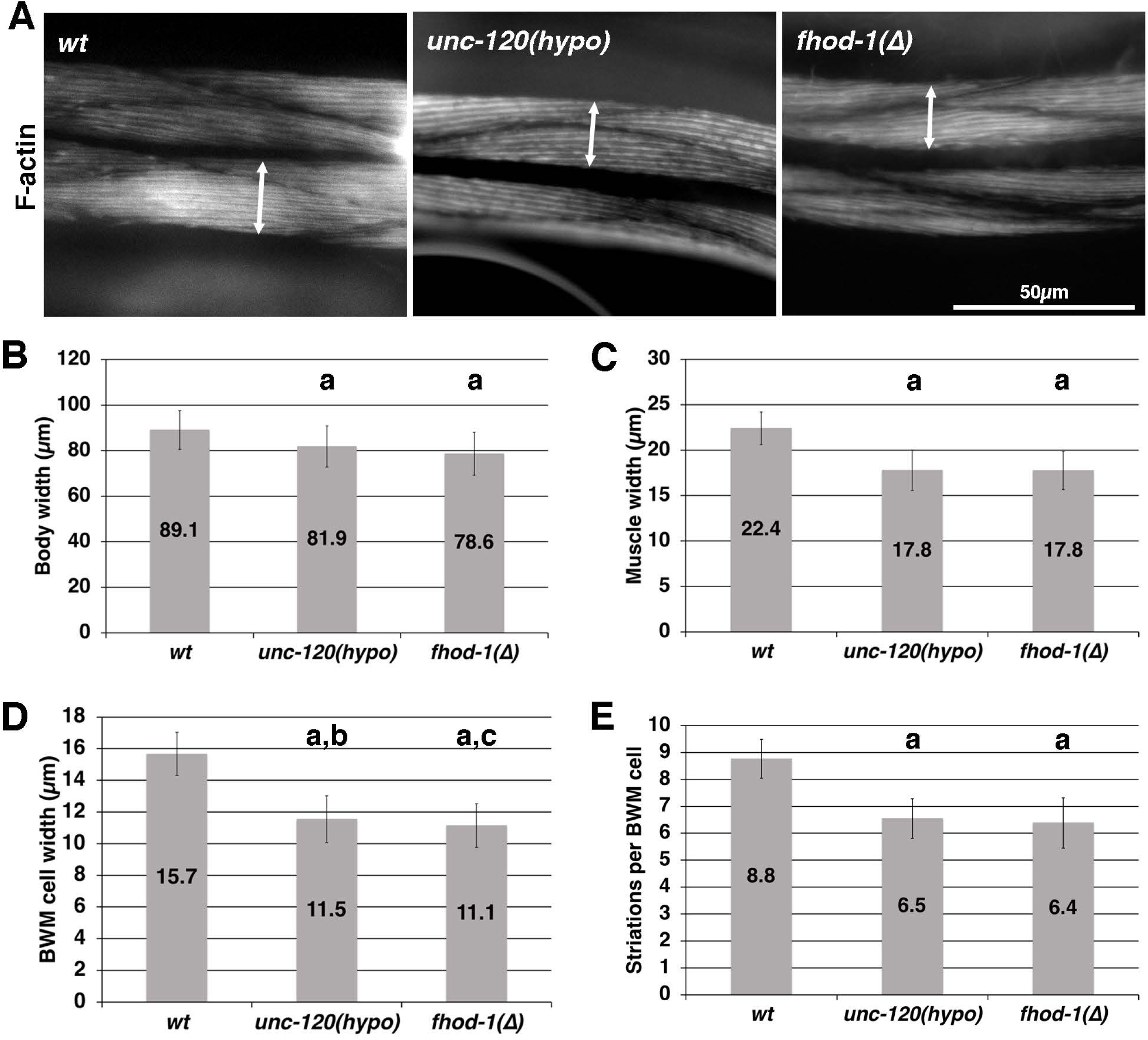
UNC-120 promotes BWM growth in a manner qualitatively similar to FHOD-1. (A) Fluorescent phalloidin stain showing dorsal views of a portion of single worms of the indicated genotypes, with two BWM quadrants extending horizontally in the field of view. Adult *unc-120(hypo)* and *fhod-1(Δ)* worms have similarly sized BWMs that are narrower than wild type (wt). Double arrows show width of one BWM. (B) Overall body widths, (C) BWM widths, and (D) individual BWM cell widths were measured. (E) The number of striations in individual BWM cells was manually counted. Shown are averages of all measurements from three independent experiments (n = 60 worms per strain, with measures of one body, two BWMs, and two BWM cells per worm). Error bars indicate one standard deviation. (a) p < 0.001 from wild type; (b) p < 0.05 from *fhod-1(Δ)*; (c) p < 0.05 from *unc-120(hypo)*. Differences in all other comparisons were not statistically significant (p > 0.05).

### Western blot analysis

For western blot analysis, samples were collected in one of two ways. By the first method, worms were washed off plates into 1.7 mL tubes and suspended in a 1:1 worm-to-M9 slurry, and sample buffer was added. Samples were boiled 3 min, disrupted with a tissue grinder 30 sec, boiled 3 min, and pelleted 15 sec. To break up genomic DNA, samples were pulled through an insulin syringe eight times before loading for gel electrophoresis. For samples prepared using this method, volumes loaded for gel electrophoresis was based on relative tubulin band intensity determined by a preliminary western blot analysis.

As an alternate method of collection, 300 worms were picked into 100 μL M9 and chilled on ice ≥ 5 min, washed once with M9, and again settled on ice 5 min. 50 μL prechilled 425-600 μm acid-washed glass beads (Sigma-Aldrich, Saint Louis, MO) were added and samples were vortexed five times 6 m/sec for 20 sec. Sample buffer was then added to samples, mixed by vortex, boiled 3 min, mixed by vortex, boiled 2 min, and pelleted 15 sec. Samples collected by this method were considered already normalized for loading for gel electrophoresis, but were prepared and analyzed in duplicate to confirm reproducibility of the normalization method and to identify uneven transfers to nitrocellulose, which were excluded from analysis. We verified that both methods of normalizing sample load sizes provided similar results for the effects of *fhod-1(Δ)* on MYO-3 levels.

Blots were probed for MYO-3 using 5-6-s diluted 1:2000, and for tubulin, if needed for loading control, using AA4.3 anti-tubulin (generated by C. Walsh, University of Pittsburgh, Pittsburgh, PA; obtained through the Developmental Studies Hybridoma Bank, University of Iowa, Iowa City, IA) at 1:1000 dilution. Goat anti-mouse conjugated to horseradish peroxidase secondary antibody (Rockland Immunochemicals, Limerick, PA) was used at 1:3000 dilution. Images were taken using ChemiDoc MP Imaging System (Bio-Rad, Hercules, CA). Band intensities were measured using Image Lab (Version 5.2.1 build 11; Bio-Rad, Hercules, CA). Excel was used for data processing. Mean and standard deviation of MYO-3 levels were determined from pooled values of all western blots after band intensities were normalized to wild type. For western blot analysis of worms treated for RNAi against *myo-3*, band intensities were normalized to the signal for the 50% dilution of control-treated wild type, because this had a more similar intensity to that for knockdown samples.

### qRT-PCR

RNA was collected following a protocol similar to Ly and colleagues [49]. Ten L2 or five L4 animals were picked into 1 µL lysis buffer (5 mM Tris-HCl, pH 8.3, 0.5% Triton X-100, 0.5% Tween 20, 0.25 mM EDTA) with 10% proteinase K (Sigma-Aldrich, Saint Louis, MO). Samples were incubated 10 min at 65°C followed by heat inactivation 1 min at 85°C. Samples were then treated 10 min with 1 U DNase I (New England BioLabs, Ipswich, MA) at 37°C, followed by 20 min heat inactivation at 75°C. cDNA was generated using Invitrogen Superscript III First-Strand Synthesis Supermix for qRT-PCR according to manufacturer’s directions (Invitrogen, Carlsbad, CA). Samples were diluted to 30 µL with diethylpyrocarbonate-treated water before use. qPCR was performed with SYBR GreenER qPCR SuperMix for iCycler Instrument (Invitrogen, Carlsbad, CA) in an iCycler (Bio-Rad, Hercules, CA) using 10 sec 55°C for annealing and 1 min 60°C for elongation. Primer sequences for *myo-3* and reference gene *ama-1,* coding for the largest subunit of RNA polymerase II, are provided in Table 1. ΔΔCt method was used for quantitation.

**Table 1.**
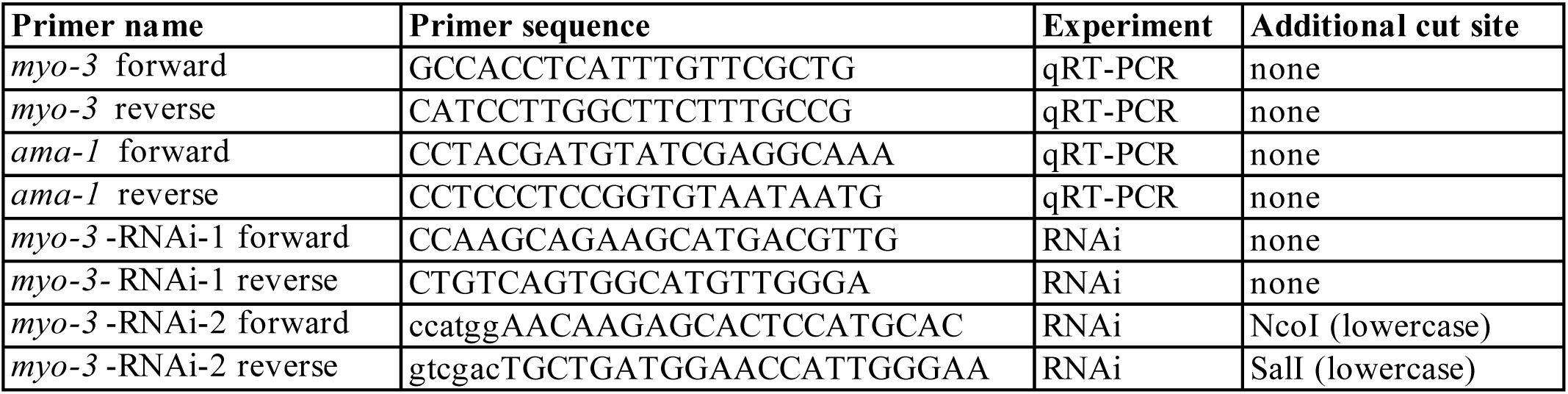
Primers used in this study. Primer names and sequences, experiments primers were used for, and identity of added restriction enzyme cut sites are indicated.

### mRNA sequencing

For each strain, four independent egg collections from bleached adult worms were used to generate four independent age-synchronized populations that were allowed to develop for 60 hours at 15°C before being collected as L3 larvae in 100 µL M9, flash frozen in liquid nitrogen, and stored at −80°C. To purify RNA, worm samples were mixed with 900 µL TRIzol reagent (Ambion, Carlsbad, CA) and vortexed 15 min before being flash frozen in liquid nitrogen. Samples were thawed at room temperature with occasional vortexing, mixed with 200 µL chloroform, and then spun at 12,000 rpm, 15 min. The aqueous layer was moved to a fresh tube, RNA was precipitated with isopropanol, and the resultant pellet was washed with 70% ethanol before being suspended in 30 µL diethylpyrocarbonate-treated water and flash frozen in liquid nitrogen. RNA samples were processed at the SUNY Molecular Analysis Core for next generation sequencing. Data were analyzed using Partek Flow (Partek, St. Louis, MO). Sequence reads were aligned with Bowtie 2 [50, 51], quantified to annotation model WBcel235-Ensembl95, and normalized by transcripts per million reads. Differentially expressed genes (defined by false discovery rate (FDR) step up ≤ 0.05, fold change > 1.5) were identified by Gene-Specific Analysis. Striated muscle genes were defined as genes contained listed in the WormBase anatomy term WBbt:0005779 [52]. To generate a heat map, Hierarchical Clustering was performed, and Venn diagrams were created using List Generator features of Partek Flow, with additional information added manually using Photoshop.

### Drug treatments

Proteolysis inhibitor dissolved in DMSO, or DMSO alone for control plates, was placed in the middle of bacterial lawns on 5 mL NGM/OP50-1 plates and allowed to diffuse overnight, to final predicted concentrations of 5 μg/mL calpain inhibitor II (Sigma-Aldrich, Saint Louis, MO) or 5 μM MG132 (EMD Millipore, Darmstadt, Germany), similar to as previously described [53]. L1 animals were washed off untreated plates and added to drug-treated plates to develop for 72 hours, before being collected as adults for western blot analysis or staining.

### RNAi knockdown

Two non-overlapping RNAi target sequences against *myo-3* were amplified by PCR from a cDNA library prepared from mixed stage wild-type animals using *myo-3* primers (Table 1). These were cloned into pCR4 Blunt-TOPO vector using Zero Blunt TOPO PCR cloning kit for sequencing (Invitrogen, Carlsbad, CA), and then transferred by standard cloning techniques into L4440, a plasmid that supports double-stranded RNA synthesis [54], to produce L4440-*myo-3*-RNAi-1 and L4440-*myo-3*-RNAi-2. These and L4440 empty vector control were transformed into HT115 bacteria. Overnight cultures of transformed bacteria were diluted 1:100 in 2xYT [47] containing 12.5 μg/ml tetracycline and 100 μg/ml ampicillin, and grown 3 hours at 37°C, then induced with 0.4 mM IPTG for 3 hours at 37°C. Induced cultures were concentrated fivefold, and 150 μL plated on RNAi feeder plates [55]. L1 stage worms were collected by washing from plates with M9, and further washed four times to remove OP50-1 bacteria before being plated on RNAi feeder plates with induced HT115 bacteria. Animals developed under RNAi conditions for 72 hours before being collected for western blot analysis, or 76 hours before being collected for phalloidin staining.

### Statistical Analyses

For comparisons of two sets of data, a student’s t-test was used to determine statistical significance. In comparisons of three or more sets of data, results were subjected to analysis of variance, followed by least significant difference *post hoc* testing. P-value < 0.05 was considered to be statistically significant.

### Data availability statement

The mRNA-sequencing data in this publication has been deposited in NCBI’s Gene Expression Omnibus [56] and are accessible through GEO Series accession number GSE158232 (https://www.ncbi.nlm.nih.gov/geo/query/acc.cgi?acc=GSE158232). Other data supporting the findings of this study are available from the corresponding author upon reasonable request.

### Ethical considerations

This study did not involve the use of human subjects or vertebrate animals. *C. elegans* is an invertebrate model that is not covered by the guidelines of National Institutes of Health definition of Laboratory Animal, and is not subject to regulation by the SUNY Upstate Medical University Institutional Animal Care and Use Committee. Sex differences in *C. elegans* are not expected to be relevant to those in humans. For that reason and for ease of animal handling, all analyses were of self-fertilizing hermaphrodite worms. All protocols and procedures were approved by the SUNY Upstate Medical University Institutional Biosafety Committee, and followed NIH Guidelines for Research Involving Recombinant DNA Molecules.

## Results

### UNC-120 (worm SRF) promotes BWM growth in a manner qualitatively similar to FHOD-1 (formin)

If the formin FHOD-1 were to promote BWM growth by stimulating SRF/UNC-120-dependent transcription of muscle genes, we might expect defects in either FHOD-1 or UNC-120 would have similar effects on BWM development. Being that a null allele of *unc-120* leads to early larval stage lethality [44], we examined the effects of the temperature-sensitive *unc-120(st364)*, called hereafter *unc-120(hypo)*, at its semi-permissive temperature 15°C. We compared BWM in these to animals bearing the deletion allele *fhod-1(tm2363)*, called hereafter *fhod-1(Δ)*, which eliminates the FHOD-1 FH2 domain and is thus presumed a null for formin-dependent actin filament assembly [40].

To visualize BWM, age-synchronized adult worms were subjected to fluorescent phalloidin staining of F-actin, which strongly stains the contractile lattice in BWM cells (Fig. 1 A). In *C. elegans*, BWMs appear as four F-actin-rich bands that reach from the head to the tail of the animal, with two BWMs positioned dorsally and two positioned ventrally. Adult BWMs are composed of flattened mononucleated BWM cells, each with a spindle-shaped contractile lattice [57]. The vast majority of BWM cell growth occurs during larval development, when the lattice widens by the addition of sarcomeres. Partial defects in BWM growth are often reflected in BWMs that are proportionately narrower compared to the overall width of the entire animal. Thus, while *unc-120(hypo)* and *fhod-1(Δ)* were somewhat smaller in body size than wild type (Fig. 1 B), their BWMs were proportionately even narrower (Fig. 1 C). Moreover, this degree of narrowing was similar in *unc-120(hypo)* and *fhod-1(Δ)* worms grown at 15°C (Fig. 1 C).

Because of the semi-staggered arrangement of BWM cells in BWM (Fig. 1 A), one might expect BWMs could narrow either due to a reduction in the number of BWM cells, or due to narrowing of the individual BWM cells. In the case of *fhod-1(Δ)*, narrow BWMs reflect a decrease in individual cell widths [40]. Similarly, we observed *unc-120(hypo)* worms had narrow BWM cells compared to wild-type worms, and this effect was also quantitatively identical to that of *fhod-1(Δ)* worms (Fig. 1 D).

Sarcomeres in the BWM cell contractile lattice are organized into oblique striations [57], such that striations are oriented at only a 5-7° angle from the longitudinal axis (Fig. 1 A). As such, the width of the BWM cell contractile lattice is essentially a function of the width of individual striations, and the total number of striations in the lattice. Therefore, BWM cell narrowing could be due to either having narrow striations or fewer striations. We previously showed that in the case of *fhod-1(Δ)*, narrowness correlates well with the presence of fewer striations per BWM cell [40], with sarcomere narrowing making a much smaller contribution [42]. Again, showing a similarity between *fhod-1* and *unc-120*, we observed *unc-120(hypo)* BWM cells had a reduction in F-actin-rich striations per BWM cell to the same degree as *fhod-1(Δ)* worms (Fig. 1 E). Thus, partial inhibition of UNC-120 activity affects BWM growth in ways very similar to those caused by absence of FHOD-1-dependent actin assembly activity, with assembly of fewer oblique striations.

### UNC-120 promotes MYO-3 protein expression similar to FHOD-1

We have previously shown that *fhod-1(Δ)* worms express less muscle-specific myosin II heavy chain MYO-3 [42]. To verify this decrease in MYO-3 is specifically due to absence of FHOD-1, we examined myosin expression in *fhod-1(Δ)* worms bearing a *fhod-1::gfp* transgene (designated *fhod-1(Δ); fhod-1::gfp*) that we had previously shown to partially rescue BWM cell growth [40]. Western blot analysis of whole worm lysates showed myosin levels were partially rescued by the transgene, confirming FHOD-1 contributes to MYO-3 protein expression (Fig. 2 A and B).

**Figure 2.**
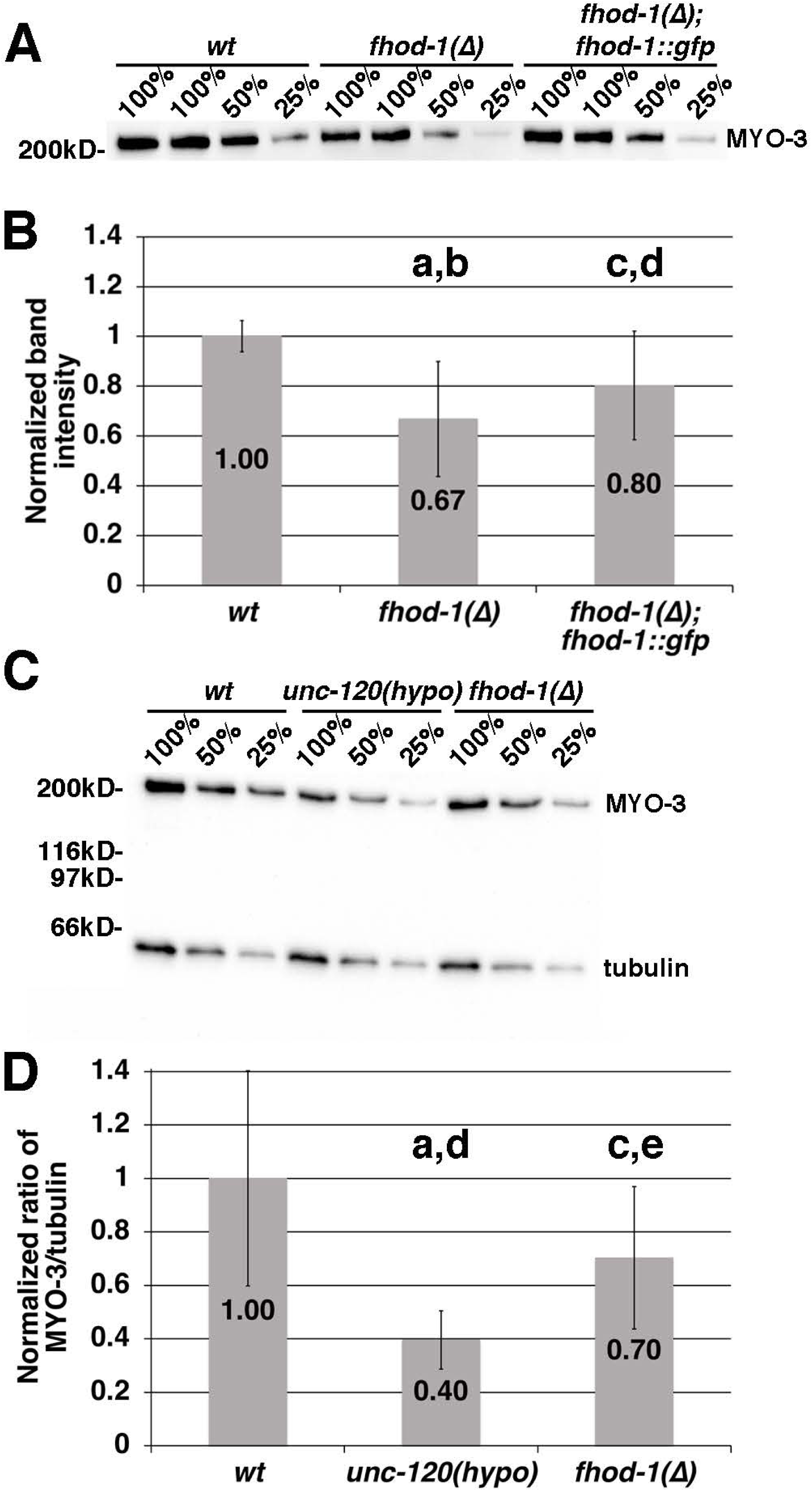
UNC-120 and FHOD-1 promote MYO-3 protein expression. (A) MYO-3 western blot of dilutions (expressed as percentages) of whole adult worm extracts show *fhod-1(Δ)* worms have less MYO-3 than wild-type worms (wt), and this is partially rescued by a *fhod-1::gfp* transgene. Sample loads were normalized based on number of worms in each sample. Position of molecular weight standard is indicated. (B) Average band intensities, from 100% and 50% samples from five independently collected lysates, were normalized to wild type, with each lysate being subjected to two independent western blots. Error bars indicate one standard deviation. (C) MYO-3 and tubulin western blots of adult worm extracts show a decrease in MYO-3 in *unc-120(hypo)* and *fhod-1(Δ)* worms compared to wild type. Sample loads were normalized based on tubulin signal. Positions of molecular weight standards are indicated. (D) Average ratios of MYO-3 to tubulin band intensities of three independently collected lysates were normalized to wild type, with each lysate being subjected to two independent western blots. Error bars indicate one standard deviation. (a) p < 0.001 from wild type; (b) p < 0.05 from *fhod-1(Δ); fhod-1::gfp*; (c) p < 0.05 from wild type; (d) p < 0.05 from *fhod-1(Δ)*; (e) p < 0.05 from *unc-120(hypo)*.

If worm FHOD-1 promotes MYO-3 expression by an SRF-dependent manner, we might expect worms partially defective for UNC-120 would have similarly reduced MYO-3 levels. Consistent with this, western blot analysis showed a decrease in MYO-3 in *unc-120(hypo)* worms that was more substantial than that in *fhod-1(Δ)* worms (Fig. 2 C and D). Thus, partial reduction of UNC-120 activity resembles absence of FHOD-1-dependent actin assembly in terms of effects on BWM cell development and on myosin expression.

### FHOD-1 does not promote transcription of muscle genes by UNC-120

If FHOD-1 were to promote muscle growth by stimulating UNC-120-dependent transcription, we would also expect *fhod-1(Δ)* and *unc-120(hypo)* worms to have similar changes in expression of muscle genes when compared to wild type. To directly test this, we performed mRNA sequencing on poly-A-enriched RNA from wild-type, *fhod-1(Δ)*, and *unc-120(hypo)* worms at the L3 larval stage, when worms are in the middle of BWM growth (Supplementary Table 1). Differentially expressed genes were defined by an FDR step up ≤ 0.05, and a fold change ≥ 1.5 (Fig. 3 A; Supplementary Table 2). Interestingly, despite *fhod-1(Δ)* and *unc-120(hypo)* having similar effects on BWM development, there were more genes differentially expressed between wild-type and *fhod-1(Δ)* animals than there were between wild-type and *unc-120(hypo)* animals (266 genes versus 73; Fig. 3 A). Moreover, only a minority of genes (32) were differentially expressed for both *fhod-1(Δ)* and *unc-120(hypo)* as compared to wild type (Fig. 3 A).

**Figure 3.**
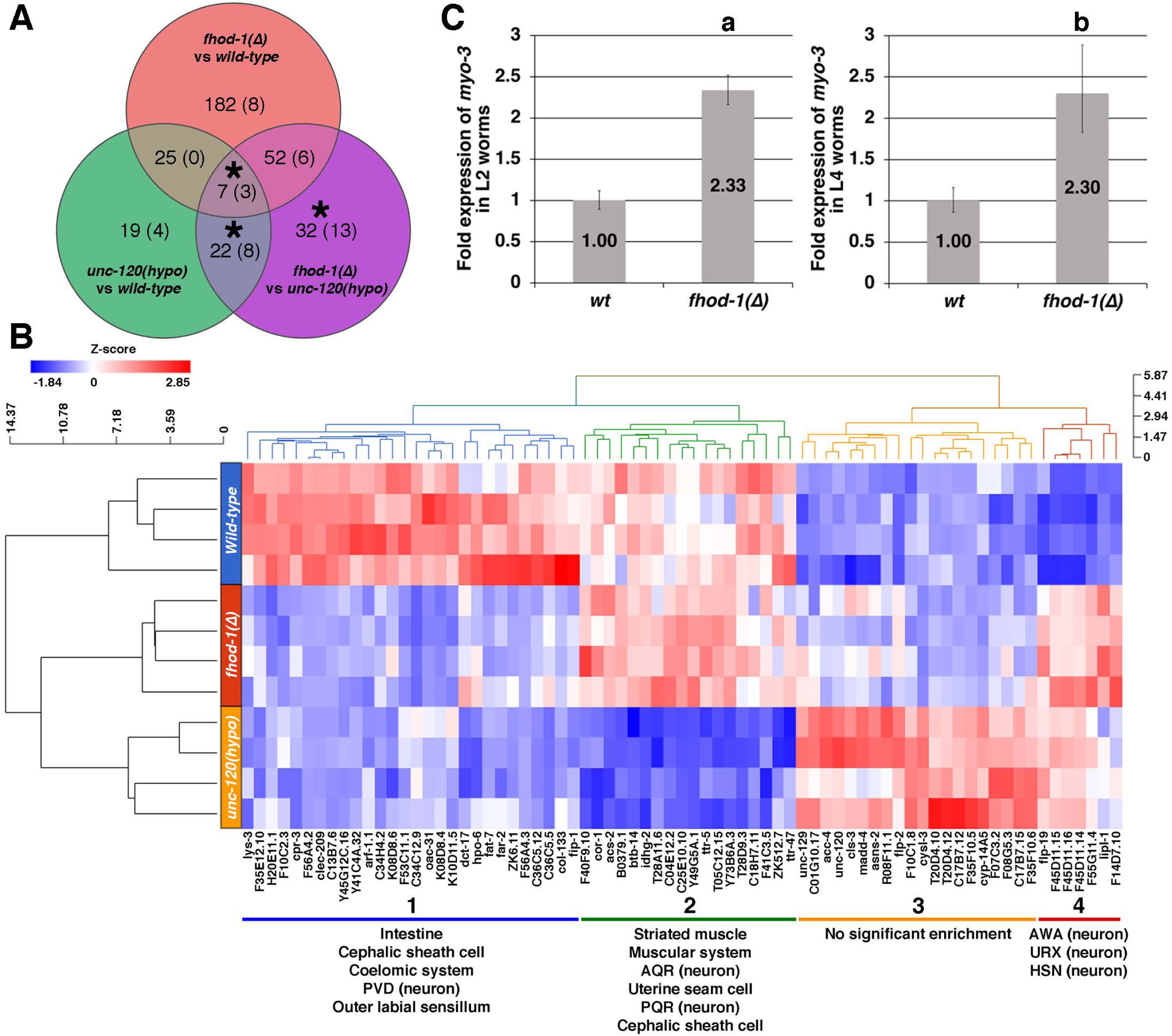
Defects in FHOD-1 and in UNC-120 have differing effects on muscle gene expression. (A) Venn diagram of differentially expressed genes (FDR step up ≤ 0.05, fold change > 1.5) identified by comparison of mRNA sequencing results for L3 stage *fhod-1(Δ)*, *unc-120(hypo)*, and wild-type worms. Indicated in each sector is the number of differentially expressed genes, and in parentheses, the number of genes expressed predominantly in striated muscle. (*) denotes significant enrichment of striated muscle genes in a gene set, based on the WormBase Enrichment Analysis Tool. Gene identities are provided in Supplementary Table 2. (B) Heat map showing the relative levels of gene expression in four replicate samples of the indicated genotypes, for all genes differentially expressed between *unc-120(hypo)* and wild-type worms. Red indicates expression was elevated above the average for the entire group, while blue indicates decreased expression. Hierarchical clustering identified four gene groups, with enrichment for tissue expression displayed beneath each group. Only group 2 was enriched for genes expressed primarily in striated muscle and the muscular system. (C) Quantitative RT-PCR values for *myo-3* were determined in comparison to reference gene *ama-1* by the ΔΔCt method, and normalized to wild type (wt). Shown are average results from three RNA samples, each analyzed in triplicate, from L2 stage and L4 stage worms. Error bars indicate one standard error. (a) p < 0.001 from wild type; (b) p < 0.05 from wild type.

Genes expressed in striated muscle were identified based on their annotation in WormBase (version WS276, anatomy term WBbt:0005779; [52]). Only 17 (6%) of genes differentially expressed between wild-type and *fhod-1(Δ)* worms are expressed predominantly in striated muscle (Fig. 3 A; Supplementary Table 2). The WormBase Enrichment Analysis Tool (version WS271) was used to determine whether gene sets were significantly enriched for expression in any particular tissue [58, 59]. This showed striated muscle genes were not significantly enriched among those differentially expressed between wild-type and *fhod-1(Δ)* worms. In contrast, striated muscle-expressed genes accounted for 15 of 73 genes (20%) differentially expressed between wild-type and *unc-120(hypo)* animals (Fig. 3 A; Supplementary Table 2), and this did represent a significant enrichment for striated muscle genes. Even more striking, 30 genes out of 113 (27%) that were differentially expressed between *fhod-1(Δ)* and *unc-120(hypo)* animals are expressed in striated muscle (Fig. 3 A; Supplementary Table 2), and this also represented a significant enrichment. Moreover, expression of 16 of those 30 striated muscle genes changed by ≥ 1.1-fold relative to wild type in opposite directions in *fhod-1(Δ)* and *unc-120(hypo)* worms, with higher expression in *fhod-1(Δ)* and lower expression in *unc-120(hypo)* (Supplementary Table 2). Another eight had similar divergent changes with higher expression in *fhod-1(Δ)* than *unc-120(hypo)* but fell within 1.1-fold of wild type for one mutant or the other (Supplementary Table 2). Of the remaining six striated muscle genes differentially expressed between *fhod-1(Δ)* and *unc-120(hypo)* worms, three were upregulated and three were downregulated in both mutants compared to wild type (Supplementary Table 2). Thus, more striated muscle genes are differentially expressed between *fhod-1(Δ)* and *unc-120(hypo)* worms than in any other strain comparison, and most of these genes changed expression in opposite directions in the two mutants.

As an alternative unbiased analysis of whether genes affected by *unc-120(hypo)* were similarly altered in *fhod-1(Δ)* worms relative to wild-type worms, hierarchical clustering was performed on all genes differentially expressed between wild-type and *unc-120(hypo)* worms (Fig. 3 B). Of four gene groups identified by this method, two (groups 1 and 4 in Fig. 3 B) showed similar changes in expression in *fhod-1(Δ)* and *unc-120(hypo)* worms relative to wild type. However, tissue enrichment analysis showed these two gene groups are enriched for expression in neurons or the intestine but not muscle. For the other two gene groups (groups 2 and 3 in Fig. 3 B), expression levels in *fhod-1(Δ)* worms were more similar to wild-type than *unc-120(hypo)* worms. Gene group 3 was not enriched for expression in any particular tissue, but gene group 2 was enriched for genes expressed in striated muscle and the muscular system. Thus, gene group 2 enriched for muscle genes has decreased expression in *unc-120(hypo)* worms, but is largely not affected in *fhod-1(Δ)* worms (Fig. 3 B).

Although *myo-3* did not meet our stringent cut-off criteria for differentially expressed genes, mRNA sequencing showed a trend of MYO-3 mRNA levels decreasing in *unc-120(hypo)* worms with a fold change −1.76, but increasing in *fhod-1(Δ)* worms with +1.13 fold expression. To verify this unexpected result, we measured MYO-3 mRNA levels in wild-type and *fhod-1(Δ)* worms by qRT-PCR, selecting two larval stages (L2 and L4) predicted to have relatively high levels of MYO-3 mRNA in wild-type worms [40, 60]. Qualitatively consistent with our mRNA sequencing results, qRT-PCR showed MYO-3 mRNA was increased in *fhod-1(Δ)* worms relative to wild type at both stages (Fig. 3 C). These results are consistent with formin/FHOD-1 affecting myosin expression through a mechanism that is distinct from SRF/UNC-120-dependent signaling.

### Proteasome-dependent degradation of MYO-3 is enhanced in *fhod-1(Δ)* worms

The overabundance of MYO-3 mRNA we observe in *fhod-1(Δ)* worms suggests defects in neither *myo-3* transcription nor MYO-3 mRNA stability cause the reduction in MYO-3 protein. This led us to examine whether there is excessive MYO-3 protein degradation in *fhod-1(Δ)* worms. Calpains have been shown to degrade a β-galactosidase reporter protein in BWM in response to several disruptions of muscle attachment structures, including dense bodies [53]. Since *fhod-1(Δ)* worms have malformed dense bodies, we examined whether enhanced calpain activity was responsible for their reduced MYO-3 levels. We treated wild-type and *fhod-1(Δ)* worms with 5 μg/mL calpain inhibitor II or DMSO vehicle control, for 72 hours (from L1 stage until adulthood), similar to what has been done previously [53]. Western blot analysis showed DMSO-treatment had no effect on MYO-3 levels, and DMSO-treated *fhod-1(Δ)* worms still had less MYO-3 than wild type (Fig. S1). Rather than observing rescue of MYO-3 expression by calpain inhibition, *fhod-1(Δ)* worms treated with calpain inhibitor II trended toward decreased MYO-3 protein compared to DMSO-treated, although this was not statistically significant, and the inhibitor had no effect on MYO-3 in wild-type worms (Fig. S1). Thus, calpains are not responsible for the reduced MYO-3 protein levels in *fhod-1(Δ)* worms.

The proteasome represents a second major protein degradation pathway in BWM [61]. Complete proteasome inactivation is lethal to *C. elegans* [62]. However, partial inhibition of the proteasome using MG132 has been shown to be effective in blocking degradation of a β-galactosidase reporter protein in BWM after genetic denervation or starvation [63]. Thus, we treated animals with DMSO or 5 μM MG132 for 72 hours (from L1 stage to adulthood), similar to as had been done previously [63]. Again, DMSO had no apparent effect on MYO-3 expression. However, MG132 treatment resulted in elevated MYO-3 in both wild-type and *fhod-1(Δ)* worms (Fig. 4). This trend was not statistically significant for wild-type worms, but MG132 treatment significantly increased MYO-3 in *fhod-1(Δ)* worms, bringing it to a level statistically indistinguishable from treated or untreated wild type (Fig. 4 B). This suggests that in the absence of FHOD-1 there is increased degradation of MYO-3 by the proteasome.

**Figure 4.**
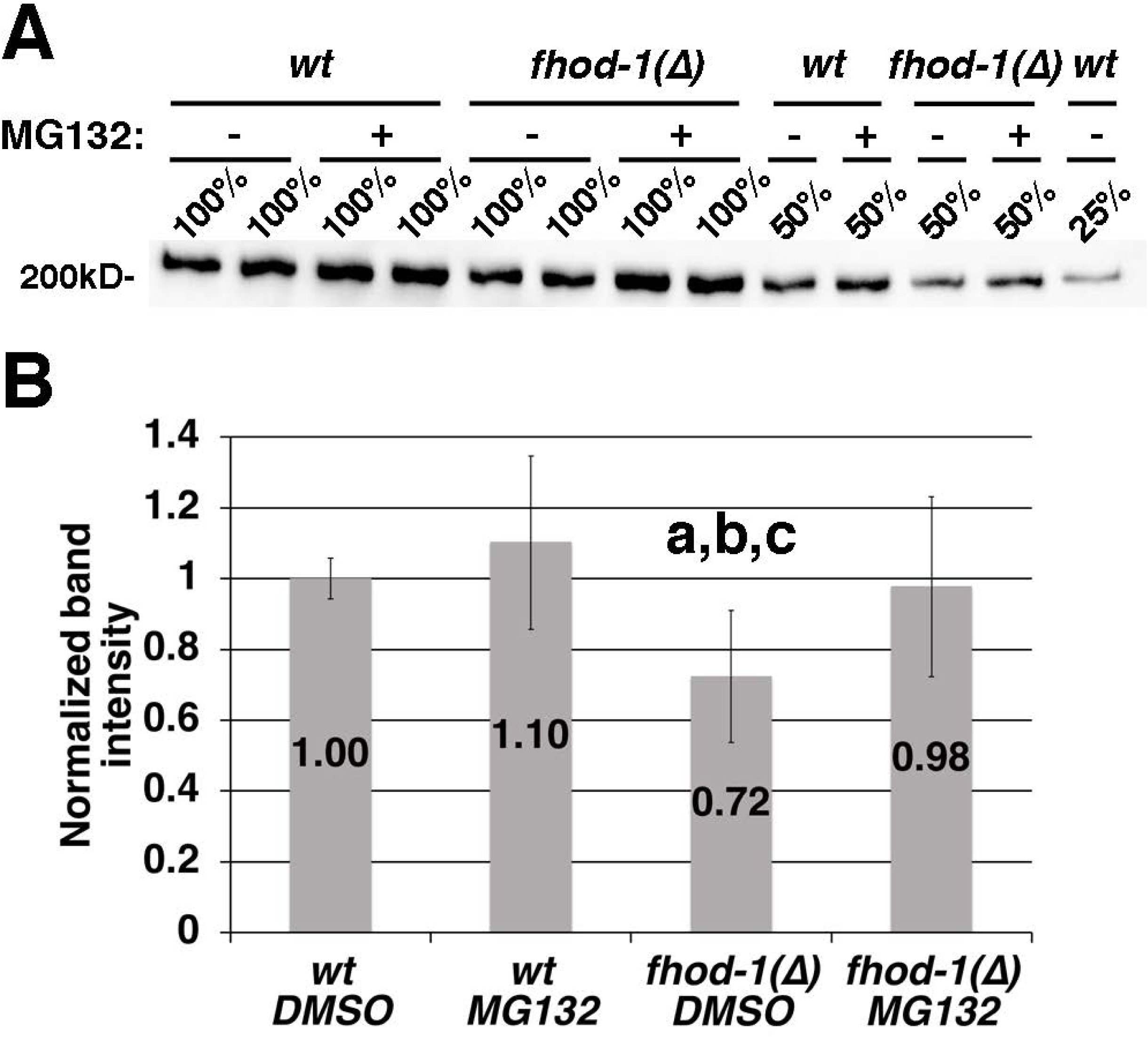
Inhibition of the proteasome restores MYO-3 to normal levels in *fhod-1(Δ)* worms. (A) MYO-3 western blot of dilutions (expressed as percentages) of whole worm extracts of adults treated with 5 μM MG132 or DMSO. Samples were prepared from equal numbers of worms. Position of molecular weight standard is indicated. (B) Shown are the averages of all measured band intensities from 100% and 50% dilutions from six independently collected lysates (normalized to wild type), with each lysate being subjected to two independent western blots. Error bars indicate one standard deviation. Treatment with 5 μM MG132 elevated MYO-3 in *fhod-1(Δ)* worms to a level statistically indistinguishable from wild-type worms (wt) treated with DMSO or 5 μM MG132. (a) p < 0.001 from wild type DMSO; (b) p < 0.001 from wild type MG132; (c) p < 0.001 from *fhod-1(Δ)* MG132. Differences in all other comparisons were not statistically significant (p > 0.05).

### Changes in MYO-3 expression are not responsible for the changes in BWM morphology in *fhod-1(Δ)* worms

Based on our evidence of elevated proteolysis, we hypothesized *fhod-1(Δ)* worms might assemble fewer sarcomeres in their BWM cells due to a deficiency in their supply of sarcomeric proteins. To test whether proteasome inhibition and restoration of wild-type MYO-3 levels were sufficient to rescue BWM development in *fhod-1(Δ)* worms, we measured the widths of BWMs and muscle cells, as well as the number of striations per muscle cell, for wild-type and *fhod-1(Δ)* worms after treatment with MG132 (Fig. 5 A-E). Fluorescent phalloidin stain of adult worms showed treatment with DMSO vehicle had no effect on any of these BWM measurements (compare Fig. 1 and Fig. 5 A-E, DMSO-treated animals). Contrary to our expectations, treatment with 5 µM MG132 also caused no change in BWM measurements (Fig. 5 A-E). Interestingly, this suggests MG132-treated *fhod-1(Δ)* worms should have an excess of MYO-3 for the number of sarcomeres present in their BWM cells. However, immunostain against MYO-3 showed the myosin still localized to the central portion of sarcomere A bands, just as it does under normal conditions [64], and we observed no aggregation or mislocalization (Fig. 5 F). Thus, restoration of MYO-3 protein is not sufficient to restore normal development to *fhod-1(Δ)* BWM cells, nor does an increase in MYO-3 levels significantly perturb muscle morphology in *fhod-1(Δ)* animals.

**Figure 5.**
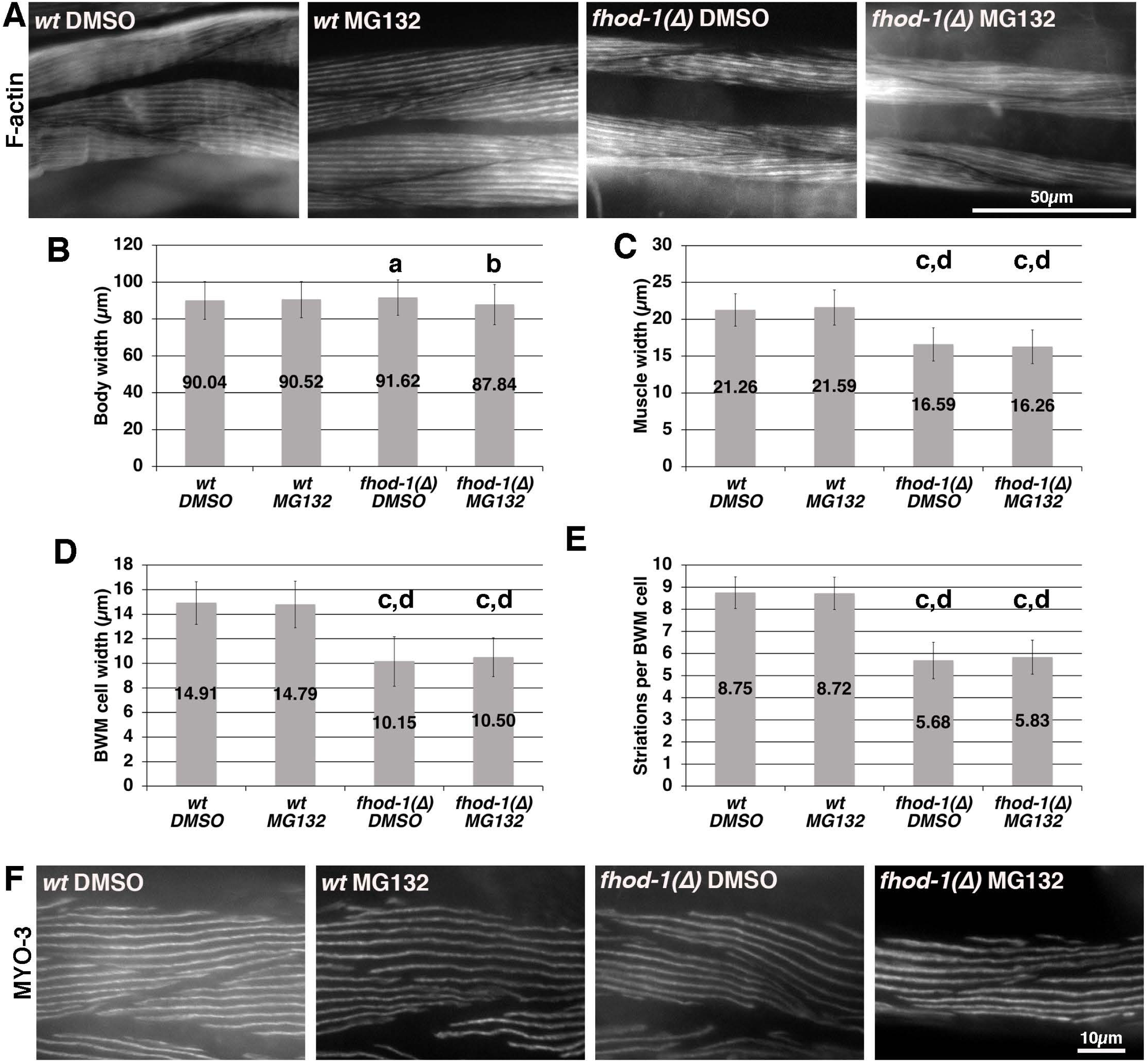
Proteasome inhibition in *fhod-1(Δ)* worms does not restore normal BWM growth. (A) Fluorescent phalloidin stain shows BWM of wild-type (wt) and *fhod-1(Δ)* adults grown in the presence of either 5 μM MG132 or DMSO. (B) Overall body width, (C) BWM widths, and (D) individual BWM cell widths were measured. (E) The number of striations in individual BWM cells was manually counted. Shown are averages of all measurements from three independent experiments (n = 60 worms per strain, with measures of one body, two BWMs, and two BWM cells per worm). Error bars indicate one standard deviation. BWM measures are identical in *fhod-1(Δ)* adults grown in the presence of either DMSO 5 μM MG132, but are consistently narrower than wild type grown under either condition. (a) p < 0.05 from *fhod-1(Δ)* MG132; (b) p < 0.05 from *fhod-1(Δ)* DMSO; (c) p < 0.001 from wild type DMSO; (d) p < 0.001 from wild type MG132. Differences in all other comparisons were not statistically significant (p > 0.05). (F) Immunostain shows MYO-3 localizes to striations in wild-type and *fhod-1(Δ)* worms, whether grown in the presence of DMSO or 5 μM MG132.

To test the converse, whether knockdown of MYO-3 would be sufficient to reduce BWM cell growth, wild-type and *fhod-1(Δ)* worms were subjected to RNAi-based knockdown of *myo-3* using two non-overlapping constructs. RNAi treatment starting in L1 larvae resulted in the reduction of MYO-3 in wild-type adults to levels comparable to or lower than control *fhod-1(Δ)* animals (Fig. S2 A and B). Despite this, we observed no effect on BWM development in wild-type or *fhod-1(Δ)* worms in response to either knockdown construct in a single test of this (Fig. S2 C-F). These results taken together suggest the reductions in BWM cell growth in *fhod-1(Δ)* animals are not driven by the changes in levels of MYO-3 protein.

## Discussion

### SRF activity is not formin-dependent in worm BWM

Many studies have shown formin-driven actin polymerization can lead to increased transcription by SRF, particularly for genes typically expressed in muscle [33–38]. For the specific case of smooth muscle genes, constitutively activated versions of the mammalian formins mDIA1, mDIA2, and FHOD1 have been shown to activate SRF-dependent transcription in multipotent 10T1/2 cells, primary aortic smooth muscles cells, and/or A7r5 smooth muscle cells [36, 37]. Conversely, expression of a dominant negative mDIA1 construct, consisting of the FH1 and FH2 domains but lacking 20 amino acids at the beginning of the FH2 domain, decreases luciferase activity driven from a known SRF target, the smooth muscle α-actin promoter, in multipotent 10T1/2 cells, primary smooth muscle cells, and A7r5 smooth muscle cells [36]. Moreover, induced overexpression of this construct in smooth muscle of adult mice led to a decrease in smooth muscle protein expression, while its constitutive overexpression in mouse smooth muscle resulted in ∼20% lethality, although the mice that survived to adulthood had no overt phenotype except delayed recovery of calponin and smooth muscle myosin heavy chain expression after arterial injury [65]. One caveat to this study, however, is that the dominant negative mDIA1 construct was described as a specific inhibitor of mDIA1- and mDIA2-dependent actin polymerization, but has been suggested by others to non-specifically inhibit actin polymerization through capping [34], leaving open the possibility it interferes with all actin polymerization, not just formin-dependent processes.

As additional evidence that endogenous formins promote transcription by SRF, knockdown of mDIA1 through RNAi in C2C12 myoblasts decreases activation of a generic SRF luciferase reporter construct [35]. Similarly, moderate (60-70%) RNAi-mediated knockdown of FHOD1 in primary smooth muscle cells reduced smooth muscle myosin heavy chain protein expression, and also blocked enhanced expression of smooth muscle myosin heavy chain, α-actin, and SM22 after sphingosine-1-phosphate treatment, a known trigger for SRF-dependent transcription [37, 66]. Together, these data suggest mDIA1, mDIA2, and FHOD1 contribute to SRF activation, at least in cultured cells. However, this still leaves open the question of whether endogenous formin activity promotes SRF activation in promoting normal muscle development *in vivo*.

We used *C. elegans* as a simple model to investigate whether formins promote transcription by SRF during striated muscle development *in vivo*. One advantage to using *C. elegans* is that FHOD-1 is the only endogenous formin that directly promotes BWM growth or sarcomere formation [41], compared to evidence that at least five formins are essential for myofibrillogenesis in cardiomyocyte cell culture [67]. Consistent with the possibility FHOD-1 and SRF might function in the same pathway in *C. elegans*, previous studies had independently shown worms bearing a *fhod-1*(*Δ*) deletion allele and worms bearing an *unc-120(hypo)* hypomorphic allele of the worm SRF gene, each exhibit thin BWMs [40, 44]. Expanding on these earlier findings, we confirmed *unc-120(hypo)* and *fhod-1(Δ)* mutant BWM cells are narrower to a quantitatively similar extent (Fig. 1). Moreover, this correlates in both mutants to fewer sarcomeres being formed per BWM cell, as opposed to the assembly of grossly smaller sarcomeres or development of fewer muscle cells per BWM, suggesting a similar mechanism. Finally, *unc-120(hypo)* and *fhod-1(Δ)* worms both have reduced amount of muscle-specific myosin II heavy chain (MYO-3), all of which would seem to support the notion that FHOD-1 and UNC-120 work together (Fig. 2).

However, despite these phenotypic similarities, we found no direct evidence FHOD-1 promotes muscle gene transcription by UNC-120. Messenger RNA sequence analysis showed *fhod-1(Δ)* perturbs expression of relatively few striated muscle genes, as compared to the effects of *unc-120(hypo)* (Fig. 3 A; Supplementary Table 2). In fact, more striated muscle genes are differentially expressed between *fhod-1(Δ)* and *unc-120(hypo)* mutants, than in comparisons of either mutant strain to wild type (Fig. 3 A; Supplementary Table 2). Also, hierarchical clustering shows a gene group enriched for striated muscle expression that is down-regulated in *unc-120(hypo)* animals is largely unaffected by *fhod-1(Δ)* (Fig. 3 B).

It is possible that this difference in the effects of worm SRF/UNC-120 and formin/FHOD-1 reflect a unique feature of *C. elegans*. The *C. elegans* gene *pqn-62* encodes a putative homolog of the myocardin/MRTF family that in mammals confers actin-sensitivity upon SRF-dependent muscle gene transcription [28, 68], and *pqn-62* is highly expressed in larval muscle [69]. However, PQN-62 has not been demonstrated to function in SRF/UNC-120 transcription or BWM development. But caveats aside, our results provide a cautionary lesson on the possibility of misinterpreting common phenotypes as evidence that formins and SRF work in a common, linear pathway in a given system.

### Formin activity is important for normal proteostasis of myosin in worm BWM

The elevated expression of a number of muscle genes in *fhod-1(Δ)* worms, including for MYO-3, led us to examine why MYO-3 protein levels are reduced in these animals, and specifically whether protein degradation was enhanced. Four main proteolytic pathways have been shown to function in worm BWM: calpains activated by disruption of integrin-based attachments (including dense bodies), the proteasome activated by starvation or denervation of muscle, lysosomes controlled by growth factor signaling, and caspases whose regulation remains unclear [61]. Of these, calpains had seemed to be the most likely mode of MYO-3 degradation based on the presence of partially disrupted dense bodies in *fhod-1(Δ)* worms [42, 53]. However, calpain inhibitor treatment did not increase MYO-3 expression in *fhod-1(Δ)* worms (Fig. S1). Instead, inhibition of the proteasome restored normal MYO-3 protein levels in *fhod-1(Δ)* worms (Fig. 4), suggesting in the absence of FHOD-1 there is increased degradation of MYO-3 (and presumably of other proteins as well) by the proteasome.

Proteasome activity has been shown to be stimulated in *C. elegans* BWM by starvation or inhibition of acetylcholine production by motor neurons [63], but it is not clear either of these effects are at play in *fhod-1(Δ)* animals. Worms lacking FHOD-1 appear well fed. We did observe transcriptional changes in neuronal genes in *fhod-1(Δ)* animals. Based on the subsets of neurons which express these genes, HSN appears to be the only motor neuron directly affected. As HSN innervates the vulval muscles to simulate egg laying in hermaphrodites [70], this might explain the reduced egg-laying capacity of *fhod-1(Δ)* animals [40, 71]. However, HSN does not synapse with BWM, and is unlikely to explain the effects of *fhod-1(Δ)* on MYO-3. It is possible FHOD-1 affects neuromuscular signaling in some other way, but we favor an alternative explanation outlined below.

The correlation between the reduction in MYO-3 levels and the reduction in muscle cell width in *fhod-1(Δ)* animals led us to question whether the *fhod-1(Δ)* muscle cell size defect was simply due to a reduced supply of sarcomeric proteins. However, whereas inhibiting the proteasome rescued expression of MYO-3 protein levels in *fhod-1(Δ)* worms, BWM growth is not rescued (Fig. 5). Conversely, knocking down expression of MYO-3 using RNAi did not affect BWM growth (Fig. S2), suggesting FHOD-1 promotes muscle growth by some mechanism other than through alteration of MYO-3 expression. As to why MYO-3 is subject to enhanced proteolysis, we hypothesize sarcomere assembly is slower in *fhod-1(Δ)* animals, and the proteasome might target the excess protein that is unable to become incorporated.

It will be interesting to see whether our finding of enhanced proteolysis by the proteasome in *fhod-1(Δ)* muscle might provide additional insight into the development of cardiomyopathy in humans. A partial disconnect is often observed between particular genetic variants and phenotypic presentation in cardiomyopathies, and it has been suggested that this may be due to factors such as modifier genes, epigenetics, environment, or incomplete penetrance [72]. Thus, mutations of the *fhod-1-*related human gene, *FHOD3*, have been suggested to cause HCM, but there is imperfect correspondence between a given *FHOD3* mutation and the development of disease [6,9–11]. Our findings point to the possibility that one contribution to variability may be interplay between *FHOD3* and perturbations in the ubiquitin proteasome system (UPS), a known contributor to cardiomyopathy [73].

FHOD3 is important for sarcomere development in the heart, and with its loss, sarcomeres fail to mature [19,21,74]. However, in terms of cardiomyopathy, it has been observed that hypertrophy-causing mutations of sarcomeric genes in general tend to be gain of function alleles, whereas loss of function mutations tend to lead to decreased contractility [75]. Extending this to FHOD3, we might anticipate HCM-associated alleles are hyperactivated, while alleles associated with DCM are hypoactive. Consistent with this, a DCM-associated FHOD3 variant is less active than wild-type FHOD3 in a luciferase-based SRF activity assay [5]. This also correlates with observations from cultured neonatal rat cardiomyocytes, where mutationally inactivated FHOD3 inhibits angiotensin II-induced hypertrophy, while mutationally activated FHOD3 leads to hypertrophy [20].

In the human heart, the UPS is the major intracellular protein degradation pathway, and is critical for normal protein turnover and function [73]. Insufficiency of the UPS is involved in cardiac hypertrophy and many cardiac diseases, including DCM. For example, E3 ligases such as muscle-atrophy F-box (Atrogin1), mouse double minute 2 (MDM2), and muscle-specific ring finger-1 (MuRF1), have been suggested to have anti-hypertrophic activity, while in DCM there is an increase in ubiquitinated proteins, suggesting decreased UPS activity [73].

Our demonstration that absence of FHOD-1 leads to an increase in proteasome activity in worm muscle suggests the possibility FHOD3 defects may also alter proteasome activity, and therefore the UPS. Somewhat similar to the case with *C. elegans* FHOD-1, absence of FHOD3 causes Z-line structures to fail to fully mature in cardiomyocytes [21, 74]. This might lead to activation of the UPS to clear unincorporated proteins, and contribute to eventual cardiac muscle failure in *FHOD3* knockout mice [21]. Conversely, *FHOD3* mutations leading to HCM might somehow reduce proteasome-dependent proteolysis of sarcomeric proteins, such as by promoting their increased incorporation into sarcomeres, thus promoting muscle hypertrophy [73]. It will be interesting to determine whether differences in intrinsic UPS activity between individuals with *FHOD3* variants contributes to the phenotypic variability found among such individuals.

## Conclusions

Overall, despite the demonstration that formin/FHOD-1 and SRF/UNC-120 both promote sarcomere assembly and myosin/MYO-3 expression in *C. elegans*, we find no evidence they function in a linear pathway. Instead, our results suggest caution should be applied when interpreting linear relationships between formins and SRF activity in muscle development based on phenotype or even protein expression levels. In *C. elegans*, formin/FHOD-1 and SRF/UNC-120 have generally opposite effects on muscle gene transcription, with loss of UNC-120 depressing muscle gene expression, while loss of FHOD-1 tends to result in unchanged or elevated muscle gene expression. Absence of FHOD-1 does lead to decreased MYO-3 protein levels, but we demonstrate this is due to elevated degradation by the proteasome. Interestingly, neither inhibition of proteasome-dependent proteolysis nor independent knockdown of MYO-3 levels are sufficient to alter muscle growth, suggesting FHOD-1 may play multiple roles in muscle, both regulating protein expression and promoting muscle growth by an as-yet unclear mechanism.

## Acknowledgements

Thanks to Frank Middleton for help with qRT-PCR and mRNA sequencing analysis, and WormBase and WormAtlas. Worm strains were obtained from the CGC, which is funded by the National Institutes of Health Office of Research Infrastructure Programs (P40 OD010440). This work was supported by the National Institutes of Health, National Institute of Arthritis and Musculoskeletal and Skin Diseases [grant number R01AR064760].

## Supplementary Table Legends

**Supplementary Table 1. mRNA sequencing data after Gene-Specific Analysis for comparisons between wild-type, *fhod-1(Δ)*, and *unc-120(hypo)* worms.** Gene symbol, total number of read counts for each gene, and p-value, FDR step up, ratio, fold change, and least-squares mean for each comparison between worm strains, for all genes following Gene-Specific Analysis.

**Supplementary Table 2. Identities of differentially expressed genes shown in Figure 3 A.** Differentially expressed genes by Venn diagram sector (Fig. 3 A), identification of genes primarily expressed in striated muscle, and fold-change of gene expression in each comparison between worm strains.

## Supplementary Figure Legends

**Supplementary Figure 1. Inhibiting calpains does not increase MYO-3 in *fhod-1(Δ)* worms.** (A) MYO-3 western blot of dilutions of whole worm extracts (expressed as percentages) prepared from equal numbers of worms treated with either 5 μg/mL calpain inhibitor II or DMSO. The position of molecular weight standard is indicated. (B) Shown are the averages of all measured band intensities from two independently collected lysates, each subjected to a single western blot. Error bars indicate one standard deviation. Treatment with 5 μg/mL calpain inhibitor II does not raise MYO-3 in *fhod-1(Δ)* worms as compared to DMSO-treated *fhod-1(Δ)* worms, and has no effect on MYO-3 in wild-type worms (wt). (a) p < 0.05 from wild type DMSO; (b) p < 0.05 from wild type calpain inhibitor II. Differences in all other comparisons were not statistically significant (p > 0.05).

**Supplementary Figure 2. Knockdown of *myo-3* after L1 stage does not change BWM growth.** (A) MYO-3 western blot of adult extracts from control worms or after treatment with two different RNAi constructs targeting *myo-3*. (B) Quantitation of measured band intensities from a single western blot shows a moderate or strong decrease in MYO-3 after RNAi treatment. (C) Overall body width, (D) BWM widths, and (E) individual BWM cell widths were measured. (F) The number of striations in individual BWM cells was manually counted. Shown are averages of all measurements from one experiment (n = 20 worms per treatment, with measures of one body, two BWMs, and two BWM cells per worm). None of these parameters appeared to change appreciably in response to *myo-3(RNAi)* treatment.

## References

[1] P.A. Harvey, L.A. Leinwand, The cell biology of disease: cellular mechanisms of cardiomyopathy, J. Cell Biol. 194 (2011) 355–365. https://doi.org/10.1083/jcb.201101100.

[2] B.J. Maron, Hypertrophic Cardiomyopathy and Other Causes of Sudden Cardiac Death in Young Competitive Athletes, with Considerations for Preparticipation Screening and Criteria for Disqualification, Cardiol. Clin. 25 (2007) 399–414. https://doi.org/https://doi.org/10.1016/j.ccl.2007.07.006.

[3] M. Kan-o, R. Takeya, K. Taniguchi, Y. Tanoue, R. Tominaga, H. Sumimoto, Expression and subcellular localization of mammalian formin Fhod3 in the embryonic and adult heart, PLoS One. 7 (2012) e34765. https://doi.org/10.1371/journal.pone.0034765.

[4] E.C. Krainer, J.L. Ouderkirk, E.W. Miller, M.R. Miller, A.T. Mersich, S.D. Blystone, The multiplicity of human formins: Expression patterns in cells and tissues, Cytoskeleton. 70 (2013) 424–438. https://doi.org/10.1002/cm.21113.

[5] T. Arimura, R. Takeya, T. Ishikawa, T. Yamano, A. Matsuo, T. Tatsumi, T. Nomura, H. Sumimoto, A. Kimura, Dilated cardiomyopathy-associated FHOD3 variant impairs the ability to induce activation of transcription factor serum response factor, Circ. J. 77 (2013) 2990–2996. https://doi.org/10.1253/circj.CJ-13-0255.

[6] E.C. Wooten, V.B. Hebl, M.J. Wolf, S.R. Greytak, N.M. Orr, I. Draper, J.E. Calvino, N.K. Kapur, M.S. Maron, I.J. Kullo, S.R. Ommen, J.M. Bos, M.J. Ackerman, G.S. Huggins, Formin homology 2 domain containing 3 variants associated with hypertrophic cardiomyopathy, Circ. Cardiovasc. Genet. 6 (2013) 10–18. https://doi.org/10.1161/CIRCGENETICS.112.965277.

[7] U. Esslinger, S. Garnier, A. Korniat, C. Proust, G. Kararigas, M. Müller-Nurasyid, J.P. Empana, M.P. Morley, C. Perret, K. Stark, A.G. Bick, S.K. Prasad, J. Kriebel, J. Li, L. Tiret, K. Strauch, D.P. O’Regan, K.B. Marguiles, J.G. Seidman, P. Boutouyrie, P. Lacolley, X. Jouven, C. Hengstenberg, M. Komajda, H. Hakonarson, R. Isnard, E. Arbustini, H. Grallert, S.A. Cook, C.E. Seidman, V. Regitz-Zagrosek, T.P. Cappola, P. Charron, F. Cambien, E. Villard, Exome-wide association study reveals novel susceptibility genes to sporadic dilated cardiomyopathy, PLoS One. 12 (2017) e0172995. https://doi.org/10.1371/journal.pone.0172995.

[8] T. Hayashi, K. Tanimoto, K. Hirayama-Yamada, E. Tsuda, M. Ayusawa, S. Nunoda, A. Hosaki, A. Kimura, Genetic background of Japanese patients with pediatric hypertrophic and restrictive cardiomyopathy, J. Hum. Genet. 63 (2018) 989–996. https://doi.org/10.1038/s10038-018-0479-y.

[9] J.P. Ochoa, M. Sabater-Molina, J.M. García-Pinilla, J. Mogensen, A. Restrepo-Córdoba, J. Palomino-Doza, E. Villacorta, M. Martinez-Moreno, J. Ramos-Maqueda, E. Zorio, M.L. Peña-Peña, P.E. García-Granja, J.F. Rodríguez-Palomares, I.J. Cárdenas-Reyes, M.M. de la Torre-Carpente, A. Bautista-Pavés, M.M. Akhtar, M.N. Cicerchia, R. Bilbao-Quesada, M.V. Mogollón-Jimenez, J. Salazar-Mendiguchía, J.M. Mesa Latorre, B. Arnaez, I. Olavarri-Miguel, M.E. Fuentes-Cañamero, A. Lamounier, J.M. Pérez Ruiz, V. Climent-Payá, I. Pérez-Sanchez, J.P. Trujillo-Quintero, L.R. Lopes, A. Repáraz-Andrade, R. Marín-Iglesias, A. Rodriguez-Vilela, M. Sandín-Fuentes, J.A. Garrote, A. Cortel-Fuster, M. Lopez-Garrido, A. Fontalba-Romero, T. Ripoll-Vera, I. Llano-Rivas, X. Fernandez-Fernandez, M. Isidoro-García, D. Garcia-Giustiniani, R. Barriales-Villa, M. Ortiz-Genga, P. García-Pavía, P.M. Elliott, J.R. Gimeno, L. Monserrat, Formin Homology 2 Domain Containing 3 (FHOD3) Is a Genetic Basis for Hypertrophic Cardiomyopathy, J. Am. Coll. Cardiol. 72 (2018) 2457–2467. https://doi.org/10.1016/j.jacc.2018.10.001.

[10] S. Huang, T. Pu, W. Wei, R. Xu, Y. Wu, Exome sequencing identifies a FHOD3 p.S527del mutation in a Chinese family with hypertrophic cardiomyopathy, J. Gene Med. 22 (2020) e3146. https://doi.org/10.1002/jgm.3146.

[11] J.P. Ochoa, L.R. Lopes, M. Perez-Barbeito, L. Cazón-Varela, M.M. de la Torre-Carpente, N. Sonicheva-Paterson, D. De Uña-Iglesias, E. Quinn, S. Kuzmina-Krutetskaya, J.A. Garrote, P.M. Elliott, L. Monserrat, Deletions of specific exons of FHOD3 detected by next-generation sequencing are associated with hypertrophic cardiomyopathy, Clin. Genet. 98 (2020) 86–90. https://doi.org/10.1111/cge.13759.

[12] D. Pruyne, M. Evangelista, C. Yang, E. Bi, S. Zigmond, A. Bretscher, C. Boone, Role of formins in actin assembly: Nucleation and barbed-end association, Science. 297 (2002) 612–615. https://doi.org/10.1126/science.1072309.

[13] I. Sagot, A.A. Rodal, J. Moseley, B.L. Goode, D. Pellman, An actin nucleation mechanism mediated by Bni1 and profilin, Nat. Cell Biol. 4 (2002) 626–631. https://doi.org/10.1038/ncb834.

[14] M. Pring, M. Evangelista, C. Boone, C. Yang, S.H. Zigmond, Mechanism of formin-induced nucleation of actin filaments, Biochemistry. 42 (2003) 486–496. https://doi.org/10.1021/bi026520j.

[15] S.H. Zigmond, M. Evangelista, C. Boone, C. Yang, A.C. Dar, F. Sicheri, J. Forkey, M. Pring, Formin Leaky Cap Allows Elongation in the Presence of Tight Capping Proteins, Curr. Biol. 13 (2003) 1820–1823. https://doi.org/10.1016/j.cub.2003.09.057.

[16] D.R. Kovar, T.D. Pollard, Insertional assembly of actin filament barbed ends in association with formins produces piconewton forces, Proc. Natl. Acad. Sci. U. S. A. 101 (2004) 14725–14730. https://doi.org/10.1073/pnas.0405902101.

[17] J.B. Moseley, I. Sagot, A.L. Manning, Y. Xu, M.J. Eck, D. Pellman, B.L. Goode, A Conserved Mechanism for Bni1- and mDia1-induced Actin Assembly and Dual Regulation of Bni1 by Bud6 and Profilin, Mol. Biol. Cell. 15 (2004) 896–907. https://doi.org/10.1091/mbc.E03-08-0621.

[18] B.L. Goode, M.J. Eck, Mechanism and Function of Formins in the Control of Actin Assembly, Annu. Rev. Biochem. 76 (2007) 593–627. https://doi.org/10.1146/annurev.biochem.75.103004.142647.

[19] K. Taniguchi, R. Takeya, S. Suetsugu, M. Kan-o, M. Narusawa, A. Shiose, R. Tominaga, H. Sumimoto, Mammalian formin Fhod3 regulates actin assembly and sarcomere organization in striated muscles, J. Biol. Chem. 284 (2009) 29873–29881. https://doi.org/10.1074/jbc.M109.059303.

[20] Q. Zhou, S.S. Wei, H. Wang, Q. Wang, W. Li, G. Li, J.W. Hou, X.M. Chen, J. Chen, W.P. Xu, Y.G. Li, Y.P. Wang, Crucial Role of ROCK2-Mediated Phosphorylation and Upregulation of FHOD3 in the Pathogenesis of Angiotensin II-Induced Cardiac Hypertrophy, Hypertension. 69 (2017) 1070–1083. https://doi.org/10.1161/HYPERTENSIONAHA.116.08662.

[21] M. Kan-O, R. Takeya, T. Abe, N. Kitajima, M. Nishida, R. Tominaga, H. Kurose, H. Sumimoto, Mammalian formin Fhod3 plays an essential role in cardiogenesis by organizing myofibrillogenesis, Biol. Open. 1 (2012) 889–896. https://doi.org/10.1242/bio.20121370.

[22] G.C.T. Pipes, E.E. Creemers, E.N. Olson, The myocardin family of transcriptional coactivators: Versatile regulators of cell growth, migration, and myogenesis, Genes Dev. 20 (2006) 1545–1556. https://doi.org/10.1101/gad.1428006.

[23] A. Parlakian, D. Tuil, G. Hamard, G. Tavernier, D. Hentzen, J.-P. Concordet, D. Paulin, Z. Li, D. Daegelen, Targeted Inactivation of Serum Response Factor in the Developing Heart Results in Myocardial Defects and Embryonic Lethality, Mol. Cell. Biol. 24 (2004) 5281– 5289. https://doi.org/10.1128/mcb.24.12.5281-5289.2004.

[24] X. Zhang, G. Azhar, J. Chai, P. Sheridan, K. Nagano, T. Brown, J. Yang, K. Khrapko, A.M. Borras, J. Lawitts, R.P. Misra, J.Y. Wei, Cardiomyopathy in transgenic mice with cardiac-specific overexpression of serum response factor, Am. J. Physiol. - Hear. Circ. Physiol. 280 (2001) H1782–H1792. https://doi.org/10.1152/ajpheart.2001.280.4.h1782.

[25] A. Parlakian, C. Charvet, B. Escoubet, M. Mericskay, J.D. Molkentin, G. Gary-Bobo, L.J. De Windt, M.A. Ludosky, D. Paulin, D. Daegelen, D. Tuil, Z. Li, Temporally controlled onset of dilated cardiomyopathy through disruption of the srf gene in adult heart, Circulation. 112 (2005) 2930–2939. https://doi.org/10.1161/CIRCULATIONAHA.105.533778.

[26] F.J. Davis, M. Gupta, S.M. Pogwizd, E. Bacha, V. Jeevanandam, M.P. Gupta, Increased expression of alternatively spliced dominant-negative isoform of SRF in human failing hearts, Am. J. Physiol. - Hear. Circ. Physiol. 282 (2002) H1521–H1533. https://doi.org/10.1152/ajpheart.00844.2001.

[27] J. Chang, L. Wei, T. Otani, K.A. Youker, M.L. Entman, R.J. Schwartz, Inhibitory cardiac transcription factor, SRF-N, is generated by caspase 3 cleavage in human heart failure and attenuated by ventricular unloading, Circulation. 108 (2003) 407–413. https://doi.org/10.1161/01.CIR.0000084502.02147.83.

[28] J.M. Miano, Role of serum response factor in the pathogenesis of disease, Lab. Investig. 90 (2010) 1274–1284. https://doi.org/10.1038/labinvest.2010.104.

[29] D. Wang, P.S. Chang, Z. Wang, L. Sutherland, J.A. Richardson, E. Small, P.A. Krieg, E.N. Olson, Activation of cardiac gene expression by myocardin, a transcriptional cofactor for serum response factor, Cell. 105 (2001) 851–862. https://doi.org/10.1016/S0092-8674(01)00404-4.

[30] J. van Tuyn, S. Knaän-Shanzer, M.J.M. van de Watering, M. de Graaf, A. van der Laarse, M.J. Schalij, E.E. van der Wall, A.A.F. de Vries, D.E. Atsma, Activation of cardiac and smooth muscle-specific genes in primary human cells after forced expression of human myocardin, Cardiovasc. Res. 67 (2005) 245–255. https://doi.org/10.1016/j.cardiores.2005.04.013.

[31] D.Z. Wang, S. Li, D. Hockemeyer, L. Sutherland, Z. Wang, G. Schratt, J.A. Richardson, A. Nordheim, E.N. Olson, Potentiation of serum response factor activity by a family of myocardin-related transcription factors, Proc. Natl. Acad. Sci. U. S. A. 99 (2002) 14855– 14860. https://doi.org/10.1073/pnas.222561499.

[32] E.N. Olson, A. Nordheim, Linking actin dynamics and gene transcription to drive cellular motile functions, Nat. Rev. Mol. Cell Biol. 11 (2010) 353–365. https://doi.org/10.1038/nrm2890.

[33] J.W. Copeland, R. Treisman, The diaphanous-related formin mDia1 controls serum response factor activity through its effects on actin polymerization, Mol. Biol. Cell. 13 (2002) 4088–4099. https://doi.org/10.1091/mbc.02-06-0092.

[34] J.W. Copeland, S.J. Copeland, R. Treisman, Homo-oligomerization is essential for F-actin assembly by the formin family FH2 domain, J. Biol. Chem. 279 (2004) 50250–50256. https://doi.org/10.1074/jbc.M404429200.

[35] S.D. Gopinath, S. Narumiya, J. Dhawan, The RhoA effector mDiaphanous regulates MyoD expression and cell cycle progression via SRF-dependent and SRF-independent pathways, J. Cell Sci. 120 (2007) 3086–3098. https://doi.org/10.1242/jcs.006619.

[36] D.P. Staus, A.L. Blaker, J.M. Taylor, C.P. Mack, Diaphanous 1 and 2 regulate smooth muscle cell differentiation by activating the myocardin-related transcription factors, Arterioscler. Thromb. Vasc. Biol. 27 (2007) 478–486. https://doi.org/10.1161/01.ATV.0000255559.77687.c1.

[37] D.P. Staus, A.L. Blaker, M.D. Medlin, J.M. Taylor, C.P. Mack, Formin homology domain-containing protein 1 regulates smooth muscle cell phenotype, Arterioscler. Thromb. Vasc. Biol. 31 (2011) 360–367. https://doi.org/10.1161/ATVBAHA.110.212993.

[38] S.F. Thurston, W.A. Kulacz, S. Shaikh, J.M. Lee, J.W. Copeland, The Ability to Induce Microtubule Acetylation Is a General Feature of Formin Proteins, PLoS One. 7 (2012) e48041. https://doi.org/10.1371/journal.pone.0048041.

[39] G.M. Benian, H.F. Epstein, *Caenorhabditis elegans* muscle: A genetic and molecular model for protein interactions in the heart, Circ. Res. 109 (2011) 1082–1095. https://doi.org/10.1161/CIRCRESAHA.110.237685.

[40] L. Mi-Mi, S. Votra, K. Kemphues, A. Bretscher, D. Pruyne, Z-line formins promote contractile lattice growth and maintenance in striated muscles of *C. elegans*, J. Cell Biol. 198 (2012) 87–102. https://doi.org/10.1083/jcb.201202053.

[41] S. Sundaramurthy, S. Votra, A. Laszlo, T. Davies, D. Pruyne, FHOD-1 is the only formin in *Caenorhabditis elegans* that promotes striated muscle growth and Z-line organization in a cell autonomous manner, BioRxiv. (2020) 2020.06.18.159582. https://doi.org/10.1101/2020.06.18.159582.

[42] L. Mi-Mi, D. Pruyne, Loss of Sarcomere-associated Formins Disrupts Z-line Organization, but does not Prevent Thin Filament Assembly in *Caenorhabditis elegans* Muscle, J. Cytol. Histol. 06 (2015) 318. https://doi.org/10.4172/2157-7099.1000318.

[43] S. Brenner, The genetics of *Caenorhabditis elegans*, Genetics. 77 (1974) 71–94.

[44] T. Fukushige, T.M. Brodigan, L.A. Schriefer, R.H. Waterston, M. Krause, Defining the transcriptional redundancy of early bodywall muscle development in *C. elegans*: Evidence for a unified theory of animal muscle development, Genes Dev. 20 (2006) 3395–3406. https://doi.org/10.1101/gad.1481706.

[45] S.G. Kuntz, B.A. Williams, P.W. Sternberg, B.J. Wold, Transcription factor redundancy and tissue-specific regulation: Evidence from functional and physical network connectivity, Genome Res. 22 (2012) 1907–1919. https://doi.org/10.1101/gr.133306.111.

[46] B.D. Williams, R.H. Waterston, Genes critical for muscle development and function in *Caenorhabditis elegans* identified through lethal mutations, J. Cell Biol. 124 (1994) 475– 490. https://doi.org/10.1083/jcb.124.4.475.

[47] K.L. Elbing, R. Brent, Recipes and Tools for Culture of *Escherichia coli*, Curr. Protoc. Mol. Biol. 125 (2019) e83. https://doi.org/10.1002/cpmb.83.

[48] M. Finney, G. Ruvkun, The *unc-86* gene product couples cell lineage and cell identity in *C. elegans*, Cell. 63 (1990) 895–905. https://doi.org/10.1016/0092-8674(90)90493-X.

[49] K. Ly, S.J. Reid, R.G. Snell, Rapid RNA analysis of individual *Caenorhabditis elegans*, MethodsX. 2 (2015) 59–63. https://doi.org/10.1016/j.mex.2015.02.002.

[50] B. Langmead, S.L. Salzberg, Fast gapped-read alignment with Bowtie 2, Nat. Methods. 9 (2012) 357–359. https://doi.org/10.1038/nmeth.1923.

[51] B. Langmead, C. Wilks, V. Antonescu, R. Charles, Scaling read aligners to hundreds of threads on general-purpose processors, Bioinformatics. 35 (2019) 421–432. https://doi.org/10.1093/bioinformatics/bty648.

[52] Wormbase WS276 (WBbt:0005779), (n.d.). http://www.wormbase.org/db/get?name=WBbt:0005779#0154--10;class=anatomy_term (accessed May 22, 2020).

[53] T. Etheridge, E.A. Oczypok, S. Lehmann, B.D. Fields, F. Shephard, L.A. Jacobson, N.J. Szewczyk, Calpains mediate integrin attachment complex maintenance of adult muscle in *Caenorhabditis elegans*, PLoS Genet. 8 (2012) e1002471. https://doi.org/10.1371/journal.pgen.1002471.

[54] L. Timmons, A. Fire, Specific interference by ingested dsRNA, Nature. 395 (1998) 854. https://doi.org/10.1038/27579.

[55] L. Timmons, D.L. Court, A. Fire, Ingestion of bacterially expressed dsRNAs can produce specific and potent genetic interference in *Caenorhabditis elegans*, Gene. 263 (2001) 103– 112. https://doi.org/10.1016/S0378-1119(00)00579-5.

[56] Edgar R, M. Domrachev, A. Lash, Gene Expression Omnibus: NCBI gene expression and hybridization array data repository, Nucleic Acids Res. 30 (2002) 207–210. https://doi.org/10.1093/nar/30.1.207.

[57] Z.F. Altun, D.H. Hall, Muscle system, somatic muscle, in: WormAtlas, 2009. https://doi.org/doi:10.3908/wormatlas.1.7.

[58] D. Angeles-Albores, R.Y.N. Lee, J. Chan, P.W. Sternberg, Tissue enrichment analysis for *C. elegans genomics*, BMC Bioinformatics. 17 (2016) 366. https://doi.org/10.1186/s12859-016-1229-9.

[59] D. Angeles-Albores, R. Lee, J. Chan, P. Sternberg, Two new functions in the WormBase Enrichment Suite, MicroPubl. Biol. 2018 (2018). https://doi.org/10.17912/W25Q2N.

[60] WormBase WS276 (myo-3), (n.d.). http://www.wormbase.org/db/get?name=WBGene00003515;class=GENE (accessed May 22, 2020).

[61] S. Lehmann, F. Shephard, L.A. Jacobson, N.J. Szewczyk, Integrated control of protein degradation in *C. elegans* muscle, Worm. 1 (2012) 141–150. https://doi.org/10.4161/worm.20465.

[62] N. Papaevgeniou, N. Chondrogianni, The ubiquitin proteasome system in *Caenorhabditis elegans* and its regulation, Redox Biol. 2 (2014) 333–347. https://doi.org/10.1016/j.redox.2014.01.007.

[63] N.J. Szewczyk, J.J. Hartman, S.J. Barmada, L.A. Jacobson, Genetic defects in acetylcholine signalling promote protein degradation in muscle cells of *Caenorhabditis elegans*, J. Cell Sci. 113 (2000) 2003–2010. http://www.ncbi.nlm.nih.gov/pubmed/10806111.

[64] D.M. Miller 3rd, I. Ortiz, G.C. Berliner, H.F. Epstein, Differential localization of two myosins within nematode thick filaments, Cell. 34 (1983) 477–490. https://doi.org/10.1016/0092-8674(83)90381-1.

[65] L. Weise-Cross, J.M. Taylor, C.P. Mack, Inhibition of Diaphanous Formin Signaling in Vivo Impairs Cardiovascular Development and Alters Smooth Muscle Cell Phenotype, Arterioscler. Thromb. Vasc. Biol. 35 (2015) 2374–2383. https://doi.org/10.1161/ATVBAHA.115.305879.

[66] K. Lockman, J.S. Hinson, M.D. Medlin, D. Morris, J.M. Taylor, C.P. Mack, Sphingosine 1-phosphate stimulates smooth muscle cell differentiation and proliferation by activating separate serum response factor co-factors, J. Biol. Chem. 279 (2004) 42422–42430. https://doi.org/10.1074/jbc.M405432200.

[67] M. Rosado, C.F. Barber, C. Berciu, S. Feldman, S.J. Birren, D. Nicastro, B.L. Goode, Critical roles for multiple formins during cardiac myofibril development and repair, Mol. Biol. Cell. 25 (2014) 811–827. https://doi.org/10.1091/mbc.E13-08-0443.

[68] WormBase WS276 (pqn-62), (n.d.). http://www.wormbase.org/db/get?name=WBGene00004145;class=Gene (accessed May 22, 2020).

[69] J. Cao, J.S. Packer, V. Ramani, D.A. Cusanovich, C. Huynh, R. Daza, X. Qiu, C. Lee, S.N. Furlan, F.J. Steemers, A. Adey, R.H. Waterston, C. Trapnell, J. Shendure, Comprehensive single-cell transcriptional profiling of a multicellular organism, Science. 357 (2017) 661–667. https://doi.org/10.1126/science.aam8940.

[70] WormAtlas (HSN), (n.d.). https://www.wormatlas.org/neurons/IndividualNeurons/HSNframeset.html (accessed May 22, 2020).

[71] A. Hegsted, F.A. Wright, S. Votra, D. Pruyne, INF2- and FHOD-related formins promote ovulation in the somatic gonad of *C. elegans*, Cytoskeleton. 73 (2016) 712–728. https://doi.org/10.1002/cm.21341.

[72] M.L. Bang, Animal Models of Congenital Cardiomyopathies Associated With Mutations in Z-Line Proteins, J. Cell. Physiol. 232 (2017) 38–52. https://doi.org/10.1002/jcp.25424.

[73] S.K. Shukla, K. Rafiq, Proteasome biology and therapeutics in cardiac diseases, Transl. Res. 205 (2019) 64–76. https://doi.org/10.1016/j.trsl.2018.09.003.

[74] A.M. Fenix, A.C. Neininger, N. Taneja, K. Hyde, M.R. Visetsouk, R.J. Garde, B. Liu, B.R. Nixon, A.E. Manalo, J.R. Becker, S.W. Crawley, D.M. Bader, M.J. Tyska, Q. Liu, J.H. Gutzman, D.T. Burnette, Muscle-specific stress fibers give rise to sarcomeres in cardiomyocytes, Elife. 7 (2018) e42144. https://doi.org/10.7554/eLife.42144.

[75] L. Mestroni, Phenotypic Heterogeneity of Sarcomeric Gene Mutations: A Matter of Gain and Loss?, J. Am. Coll. Cardiol. 54 (2009) 343–345. https://doi.org/10.1016/j.jacc.2009.04.029.

