## Supplementary figures and images for "FHOD formin and SRF promote striated muscle development through separate pathways in *C. elegans*"

### Fig. S1

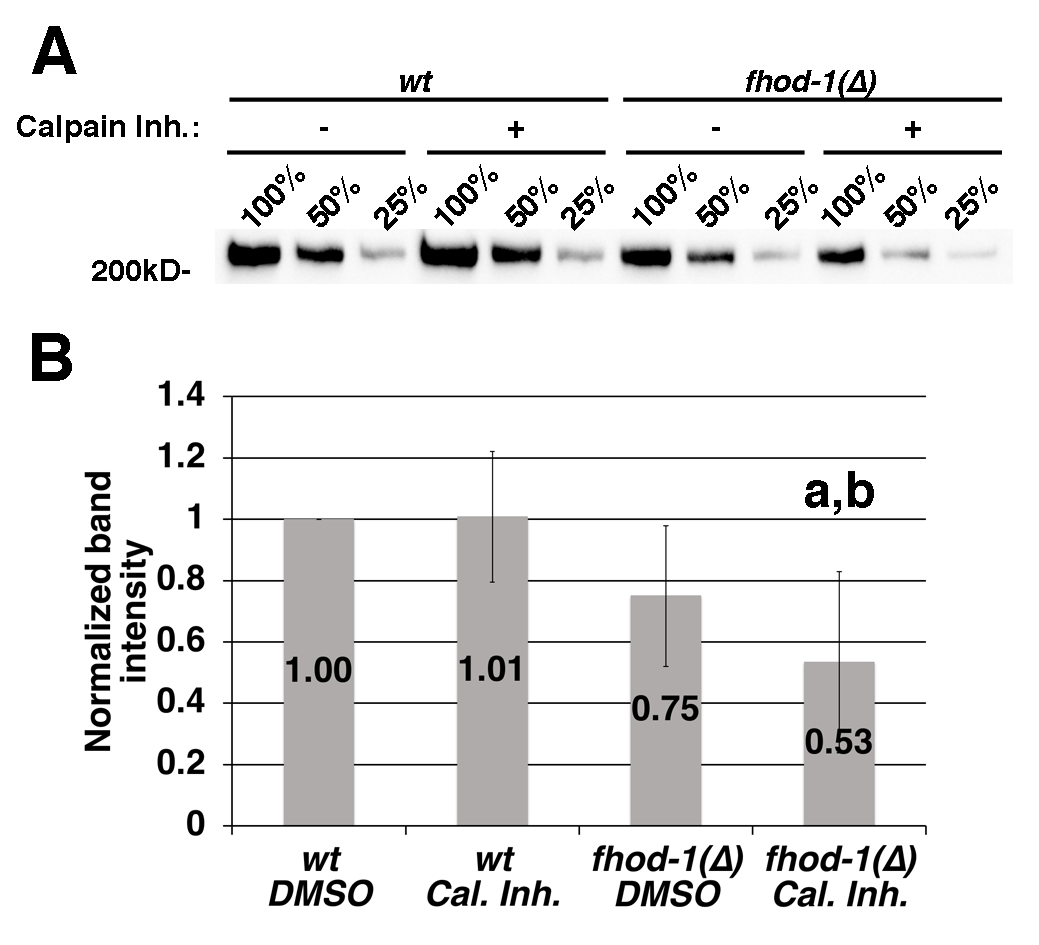

### Fig. S2

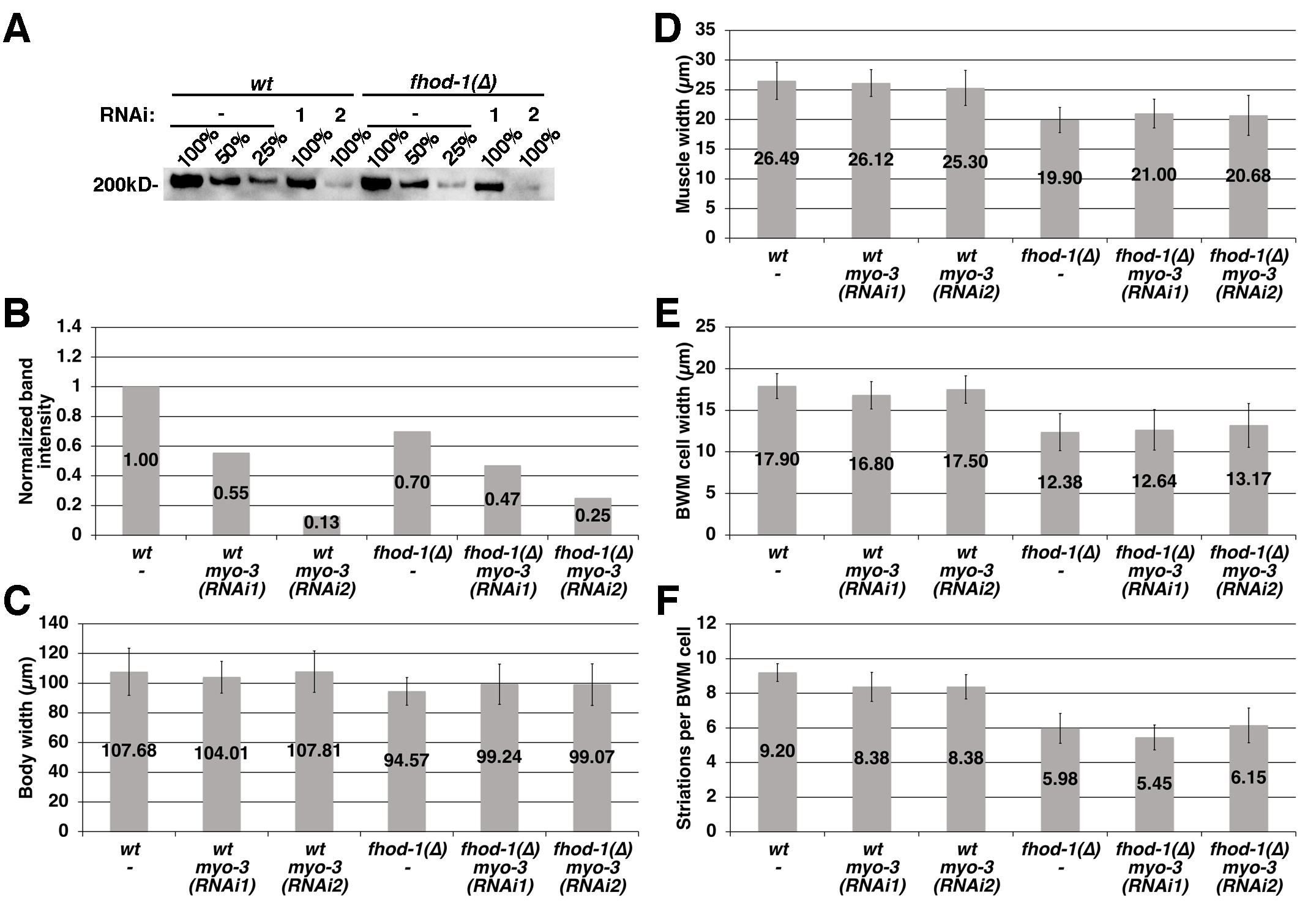
